# Probing the neural geometry of conscious object perception in time

**DOI:** 10.64898/2026.10.13.758543

**Authors:** Ayaka Hachisuka, Thomas J. Baumgarten, Dillan Spector, Lua Koenig, Biyu J. He

## Abstract

Conscious object recognition is a critical brain function, key to making sense of the plethora of visual inputs and guiding adaptive behavior. Yet, the sequence of neural processing underlying conscious object recognition remains unclear. Existing proposals alternatively suggest that conscious object recognition proceeds in a “coarse-to-fine” sequence or a “fine-to-coarse” sequence. Here, we recorded high-density EEG while participants of both sexes performed a threshold-level object recognition task under different viewing conditions. Both superordinate and basic-level category information emerged early in neural activity, followed by a late superordinate category representation in an abstract format that is disentangled from low-level object features. Strikingly, in the early period, while basic-level category information positively correlated with perceptual awareness, superordinate category information negatively predicted perceptual awareness. By contrast, both superordinate and basic-level category information positively correlated with objective categorization accuracy in the early period. In the late period, both levels of category information positively correlated with awareness. These results suggest that neural processing underlying conscious object perception proceeds from the basic to the superordinate category level.

**Significance statement:** Despite significant interests, the sequence of neural processing underlying conscious perception remains unclear. Here, we investigated the unfolding of neural events in time that support perceptual awareness of visual objects. Using a threshold-level object perception task involving three image conditions (normal, low-pass filtered, masked) and high-density EEG, we reveal that basic and superordinate category information both emerge early in the neural activity (at ∼100 ms), but only the basic-level category representation positively contributes to awareness. By contrast, the early superordinate category representation negatively correlates with perceptual awareness. At ∼300 ms, an abstract superordinate category representation emerges, which also positively correlates with perceptual awareness. Therefore, category-level neural processing underlying conscious object perception proceeds from the basic to the superordinate level.

## INTRODUCTION

Subjectively, visual object perception appears to happen instantaneously. However, object perception is supported by a hierarchical machinery with neurons at different stages of the cortical hierarchy representing visual features with different levels of detail and abstraction: from colors and lines (e.g., brown and straight) to basic object categories (e.g., a chair) to superordinate, abstract categories (e.g., furniture). Although a wealth of evidence suggests the importance of both feedforward and feedback processing in object recognition (DiCarlo et al., 2012; Mashour et al., 2020; He, 2023), currently, the sequence of processing directly underlying our perceptual awareness of an object remains unknown.

A prominent theory (Hochstein, 2002) suggests that conscious vision proceeds in a coarse-to-fine sequence: “conscious perception begins at the hierarchy’s top, gradually turning downward as needed.” Under this framework, the feedforward pathway is unconscious, and conscious perception corresponds to a feedback wave of activity. This idea resonates with the Recurrent Processing Theory (Lamme, 2006), which postulates that re-entrant (i.e., feedback) activity yields conscious experiences. Supporting the coarse-to-fine theory, behavioral studies have shown that human subjects have a behavioral advantage in reporting the global shape, as compared to the local shape, of abstract patterns (Campana et al., 2016), and that subjects can discern the category of naturalistic scenes with very brief presentation times (Fei-Fei et al., 2007). In visual search paradigms, the ‘pop out’ effect of a visual target amongst distractor stimuli appears to rely on high-level vision instead of low-level vision (Hochstein, 2002). In addition, a recent study investigating face processing reported that gender and age information emerges earlier than identity information in neural activity, suggesting that the “coarse-to-fine” sequence might also apply to a specific type of stimuli (Dobs et al., 2019). However, these studies only assessed the objective behavioral performance, such as discrimination accuracy and reaction times, or the decoding of stimulus information from neural activity, without directly measuring perceptual awareness. As a result, the identified coarse-to-fine sequence of processing might be confounded by unconscious processing, which can substantially dissociate from conscious processing (Ben-Haim et al., 2021).

On the other hand, a long-standing line of behavioral work supports a “basic category advantage” in object perception. These studies showed that participants spontaneously name objects at the basic level before the superordinate level (Rosch et al., 1976; Jolicoeur et al., 1984), that basic-level categorization is above the chance-level as soon as the object is detected (Grill-Spector and Kanwisher, 2005), and that participants can accurately identify the basic-level category with a mere 13-ms presentation of each image in a rapid-serial-visual-presentation (RSVP) task (Potter et al., 2014). Because the basic category level (e.g., dog) is more fine-grained than the superordinate category level (e.g., animal), these findings appear to support a fine-to-coarse sequence of processing during object recognition. Invasive recordings from the temporal lobe in humans (Liu et al., 2009) and non-human primates (Hung CP, 2005) also support a fast extraction of basic-level category information, at ∼100 ms after stimulus onset. However, most of these studies also only probed the objective discrimination performance or neural decoding without assessing perceptual awareness. Thus, they also do not reveal the sequence of neural processing directly underlying conscious perception. Underscoring this point, a recent fMRI study reported widespread object category representations in the brain even on unconscious trials (Mei et al., 2022).

Therefore, to test the coarse-to-fine processing theory of conscious perception and to elucidate the sequence of neural processing that directly underlies conscious object recognition, it is crucial to manipulate and measure awareness. In this study, we fill this gap by investigating the sequence of category-level neural processing that directly contributes to the perceptual awareness of an object image.

To this end, we developed a threshold-level object recognition task to measure the superordinate and basic-level categorization performances across varying levels of conscious awareness. In different conditions, we presented the original object images, images that are low-pass filtered to degrade feedforward processing, or visually masked images that disrupt recurrent processing. Recording neural activity with high-density EEG, we found that both basic-level and superordinate category information emerges relatively early (at ∼100 ms) in neural activity. However, only the early basic-level category information positively correlated with perceptual awareness ratings, whereas the early superordinate category representation *negatively* predicted perceptual awareness ratings. At a later stage, an abstract superordinate category representation emerged, which also positively correlated with perceptual awareness. By contrast, both types of category information positively correlated with the objective categorization performance. Together, these findings suggest that although an early superordinate category representation exists and supports categorization behavior, it is largely unconscious, and that category-level neural processing underlying conscious object perception proceeds from the basic to the superordinate level.

## METHODS

### Participants

39 participants (20 male, 19 female, mean age: 27 years) were recruited from the greater New York City area. All participants provided written informed consent, and the experiment was approved by the Institutional Review Board of New York University Grossman School of Medicine (protocol #s15-01323).

All participants had normal or corrected-to-normal vision and were right-handed. EEG data from 6 participants were not collected due to poor staircase behavior (based on failure to reach a constant contrast threshold for 3 or more images), 1 participant had a missing trigger file, and data from 2 participants could not be used due to an intermittently malfunctioning button response box. Thus, data from 30 participants (14 male, 16 female, mean age: 26 years) were analyzed for the main behavior and EEG analyses.

### Task design

The task was created using E-Prime 2.0 and presented on a 1920 x 1080 monitor with a 120 Hz refresh rate. The monitor was placed 63 cm away from the participant’s eyes. Participants placed their heads on a chin rest to minimize head movements and maintain a consistent viewing angle.

#### Stimuli

Our stimuli set consisted of two superordinate domains (animal/vehicle), 2 basic categories per domain (dogs/cats for animal and cars/trucks for vehicle), and 3 exemplar images per basic category, for a total of 12 exemplar images. Images were selected from the public domain, matching object position/pose/orientation between the two basic categories within each superordinate domain. Images were cropped to 250 x 250 pixels, and the resolution was adjusted to 96 dpi. Luminance was normalized using *histmatch* from the SHINE toolbox (Willenbockel et al., 2010). Additionally, the amplitude spectrum was adjusted with SHINE *sfMatch*, such that the average spatial frequency content was equated across all image stimuli.

Stimuli size was 2.4 dva, and all targets appeared pseudo-randomly at one of four locations in the periphery at 3.2 dva from fixation. The 4 dots around the target were each 1.2 dva in diameter and each dot was placed at 0.27 dva from a corner of the target image. The remaining three locations had phase-scrambled images created from original images that fell in the other three basic categories as compared to the target image (thus, if the target is a cat image, the simultaneously presented phase-scrambled images were created from a dog image, a truck image, and a car image, respectively). The overall luminance, contrast, and spatial frequency distributions were matched between target images and phase-scrambled images. Presenting stimuli in the periphery increased task difficulty to mitigate behavior ceiling effects and to induce the Object Substitution Masking effect (Koivisto et al., 2014).

#### Image conditions

Image conditions consisted of: Normal (unaltered images), Masked (through an Object Substitution Orientation-Shifted (OSOM) mask; behavioral piloting results are described in **Supplementary Results, Supplementary Methods**, and **Fig. S1**), and LSF (low spatial frequency images). For the LSF manipulation, images were low-pass filtered in the Fourier domain using a Gaussian filter. The Fourier transform of the image was multiplied by a circular mask with a cutoff radius of 12 pixels, and the filtered image was obtained via inverse Fourier transform. On each trial, one of the three image conditions was chosen in a pseudorandom manner. For all images, regardless of image manipulation, the 4-dot mask appeared with the target image for the entire duration of the 60 ms image presentation; one at each corner as described in the previous section. The remaining three locations had phase-scrambled images and did not have dots in the corner. In the Normal and LSF conditions, the presentation of the target image and surrounding 4-dots was followed by a 300 ms blank screen; only the fixation dot remained on the screen. In the Masked condition, the target image presentation was followed by a 300 ms presentation of the mask (90°-rotated version of the 4 dots that initially appeared with the target image) instead of the blank screen.

#### Staircase

Before starting the main task, contrast levels were titrated for each exemplar image (for a total of 12 interleaved staircases) in the Normal condition using an adaptive staircase procedure, to reach a ∼63% basic categorization performance for each exemplar. From a starting value of 100% contrast, each staircase began with an initially large step-size of a 45% decrease in contrast and then followed a 3-down/1-up rule to converge on the target performance level. The step sizes were reduced progressively after a certain number of reversals: from 45% to 15% after 2 reversals, to 9% after 4 reversals, and to 3% after 6 reversals. The 1-up/3-down rule meant that increases in contrast were always 1/3 of the decreases in contrast, such that the staircase converged on an approximate target of 63% performance. A maximum of 10 reversals were shown after which the staircase procedure terminated, and the final contrast values were computed as the average of the last 6 reversals. During the staircase, the target image was randomly presented in one of four locations, similar to the main task. Participants who failed to reach a stable contrast threshold for 3 or more exemplar images were rejected from the study (n = 6).

#### Main task

Each session consisted of a total of 864 trials, evenly distributed across 288 trials per image condition, 144 trials per superordinate category (animal, vehicle), 72 trials per basic level category (dog, cat, car, truck), 24 trials per exemplar, and 6 per exemplar, per position. In each session, trials were subdivided into 6 blocks, with 144 trials per block and 1 specific exemplar/position per block.

Each trial began with a 500 ms presentation of a white dot, followed by a 400-600 ms jittered fixation period (shuffled discrete values of 400/500/600 ms for the first three subjects, and taken from a uniform distribution with the same range, for the rest of the subjects), during which the fixation dot turned black. Next, the target image was presented for 60 ms at one of the four peripheral locations, surrounded by four white dots at each corner, along with scrambled images in the other three peripheral locations.

Then, two response prompts were shown: the first response screen asked, “Animal or Vehicle?” with a 2AFC task. If the participant chose “Animal,” the next prompt asked, “Cat or Dog?” If they chose “Vehicle,” the next prompt asked, “Car or Truck?” Thus, the second prompt depended on the participant’s response to the first question, regardless of whether they were correct or not. The button-response mappings were randomized across trials. Lastly, participants were asked to rate their subjective experience of vividness using the perceptual awareness scale, ranging from 0 (“I did not see the target image on the screen, not even the faintest sensation”), 1 (“I caught a glimpse of ‘something’ there, but I’m not sure what I saw”), 2 (“I got a sense of what the presented stimulus might be, but I’m not sure”), and 3 (“I saw the stimulus clearly, without any doubt”). All response screens had a maximum time limit of 2 seconds; if the participant did not respond within this time, the task proceeded to the next screen.

We adopted the above order of response prompts for two reasons. First, for the category questions, this order allows the basic-level prompt to be constrained by the superordinate response, such that both categorization questions follow a 2AFC format, resulting in comparable cognitive demand (i.e., choosing between 2 options) for both questions. Second, the probe about objective performance (categorization) preceded the probe about subjective awareness (PAS rating) (Ramsoy and Overgaard, 2004), following an established approach in the field where both performance and awareness are probed on each trial (Del Cul et al., 2007; Li et al., 2014; Levinson et al., 2021; Lanfranco et al., 2024). This task format encourages genuine guessing on the 2AFC questions when the awareness is low (because subjects may be reluctant to guess if they had answered “unseen” first), allowing better capture of unconscious processing.

### Behavioral analysis

A two-way repeated measures ANOVA was conducted using Jeffrey’s Amazing Statistics Program (JASP) (Team, 2024). For categorization performance analyses, the ANOVA was applied to the behavioral data for each participant with the following factors: 1) category level (superordinate vs. basic) and 2) image condition type (Normal, Masked, and LSF). For all behavioral analyses, the basic-level categorization performance was computed only on correct superordinate categorization trials (where valid options are presented), while superordinate categorization performance was quantified using all trials. From the “afex” package in R, the Q-Q plots of the model residuals appeared approximately linear, and the Shapiro-Wilk test did not indicate a significant deviation from normality (Shapiro-Wilk test, W = 0.981, p = 0.28). For the assessment of PAS ratings across image conditions, a one-way repeated measures ANOVA was instead applied to the mean PAS rating for each participant with the image condition (Normal, Masked, and LSF) as a factor. Again, the Q-Q plots of the model residuals were approximately linear, and the Shapiro-Wilk test did not indicate a significant deviation from normality (Shapiro-Wilk test, W = 0.986, p = 0.79). Then, for both analyses, post-hoc comparisons were applied to examine the significant main effects. P-values for post-hoc analyses were adjusted using the Bonferroni correction for multiple comparisons.

To implement a 3-way repeated measures ANOVA to assess the superordinate and basic level categorization performances across the 4 PAS ratings, the “afex” package in R was used. As in the previous section, basic-level categorization performance was quantified using correct superordinate categorization trials, while superordinate categorization performance was quantified using all trials. Within-subject factors included 1) category level (superordinate vs. basic), 2) image condition (Normal, Masked, and LSF), and 3) PAS ratings (0-3). The dependent variable was the categorization performance (with 50% as the chance level). For effects involving factors with more than two levels, Greenhouse-Geisser-corrected degrees of freedom are reported. The Shapiro-Wilk test on the model residuals indicated a significant deviation from normality (W = 0.925, p = 2.12e-11); however, given the large sample size and robustness of repeated-measures ANOVA to non-normality (Glass et al., 1972), the analysis was still performed. To assess the robustness of the results, a non-parametric permutation-based repeated measures ANOVA was conducted using the “permuco” package in R. Model residuals were permuted 10,000 times within participants to generate a null distribution of F-statistics, against which the observed F-statistics were compared. This analysis yielded similar results to the ANOVA (as reported in the Results section). Additionally, the main effect of image condition was significant (F_1,2_ = 4.27, p = 1.87e-2), consistent with the two-way ANOVA results. Post-hoc pairwise comparisons were performed using the estimated marginal means with the “emmeans” package in R to compare the superordinate and basic categorization performances at each PAS rating. P-values for post-hoc analyses were adjusted using the Bonferroni correction for multiple comparisons.

### EEG recording

Participants performed the task under EEG recordings inside a dimly lit, soundproof, electromagnetic interference-shielded room. Continuous EEG data were collected using a 128-channel BrainVision actiCAP EEG system (Brain Products GmbH). Electrodes were placed according to the international 10-20 system, and high-chloride, abrasive electrolyte gel (Abralyt HiCl, EasyCap) was used. Two mastoids were used as reference, and 4 EOG electrodes were placed around the eye (two at the outer corner of each eye, and two above and below the left eye) to capture blinks and saccades. Data was collected in DC recording mode using the BrainAmpDC system (Brain Products GmbH) with a sampling rate of 1000 Hz.

### EEG preprocessing

EEG data were preprocessed using FieldTrip (Oostenveld et al., 2011) and custom scripts in MATLAB. Continuous EEG and EOG data were bandpass filtered between 0.05 and 150 Hz and then notch filtered at 59–61 Hz and 119-121 Hz. All filters were applied offline using a symmetrical Butterworth filter with zero phase shift. Then, jump and muscle artifact rejection were semi-automatically performed. Independent component analysis (ICA) was then applied to all EEG and EOG electrodes using data filtered at >1 Hz. (Using a relatively aggressive filter prior to ICA has been found to improve the detection of eye movement artifacts (Castellanos and Makarov, 2006).) ICA components were manually inspected and noisy components were identified, including blinking, eye movement, heart rate, and muscle artifacts. Finally, the resulting weight matrices of the manually identified noisy ICs were applied to the original data (bandpass filtered at 0.05–150 Hz) for artifact rejection.

In addition, a trial-by-trial manual inspection was performed; trials that still had major artifacts were manually removed. Exceptionally noisy channels were also removed by manual inspection and then interpolated based on the mean of neighboring channels. Then, the data were epoched from -0.4 to 0.8 s, low-pass filtered at 35 Hz, and baseline-corrected by subtracting the mean of the pre-stimulus period, from -0.4 to 0 s.

### Multivariate Pattern Analysis

Time-resolved multivariate pattern analyses in Fig. 3 were conducted for each participant using the CoSMoMVPA toolbox. For each time point, voltage values from all EEG channels are input into the classifier using a cross-validation scheme described below. Linear discriminant analysis (LDA) classifiers were fit separately for each category level, as illustrated in Table 1: for the superordinate category: (1) animal vs. vehicle; for the basic-level category: (2a) dog vs. cat, (2b) car vs. truck; and for the exemplar image: (3a) dog 1 vs. dog 2 vs. dog 3 using a pairwise decoding approach, (3b) cat 1 vs. cat 2 vs. cat 3, (3c) car 1 vs. car 2 vs. car 3, (3d) truck 1 vs. truck 2 vs. truck 3.

**Table 1:**
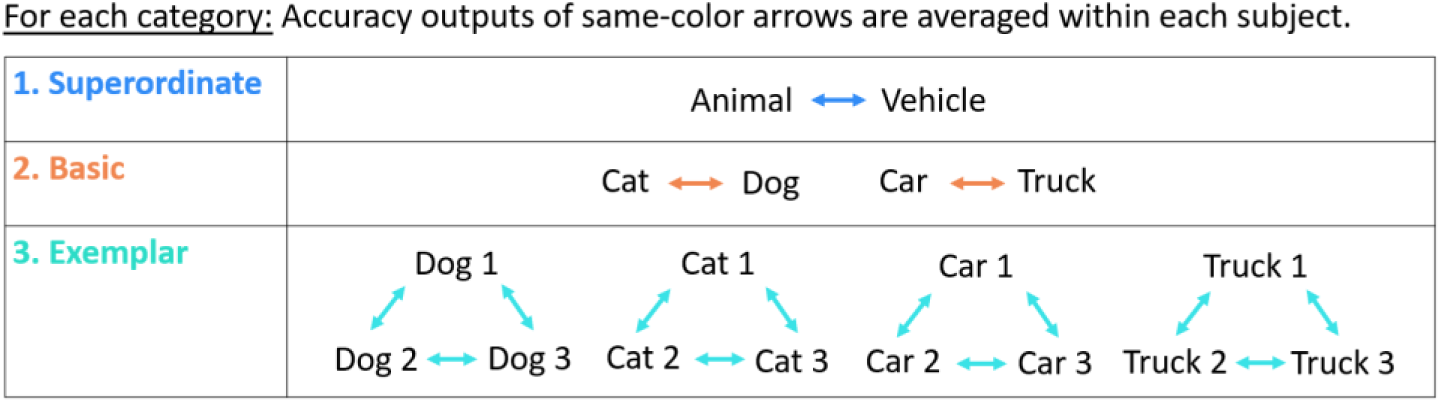
Decoding analysis schematic for superordinate, basic-level, and exemplar category decoding analysis (corresponding to Fig. 3 and Fig. S3).

To avoid overfitting, analyses were performed using a stratified cross-validation procedure with 300 iterations. Within each iteration, trials were first balanced within each participant’s dataset by randomly subsampling to match the minimum class count. Importantly, balancing was performed at the exemplar level, not just at the superordinate (equating animal and vehicle trials) or basic (equating dog and cat trials, or car and truck trials) category levels. For example, in the basic-level classifier between dogs and cats, trials were balanced such that there were equal numbers of each of the three dog types (e.g., dog 1, dog 2, and dog 3) and each of the three cat types (e.g., cat 1, cat 2, and cat 3). This ensured equal representation of all exemplar types and controlled for variability associated with not only category identity but also exemplar identity. Exemplar-level decoding was performed using a one-vs-one classification scheme within each basic level category type, with exemplar classes balanced by randomly subsampling trials to equalize exemplar (class) counts. Then, 90% training and 10% testing partitions were created by assigning approximately 10% of the total trials to the test set in multiples of the full exemplar set (i.e., in multiples of all 12 exemplar types for the superordinate category, or 6 total exemplars of dogs and cats, or 6 total exemplars of cars and trucks, for the basic level category), ensuring that the training set retained balanced exemplar counts.

After the train-test partitions were created, trials in the training set were averaged over randomly chosen groups of 4 trials to increase the signal-to-noise ratio prior to classifier training (Grootswagers et al., 2017). This step was skipped in analyses involving “undetected” trials (with PAS = 0, as in Fig. 3G and Fig. S4A-B) due to insufficient trial counts. Then, within each training set, the Linear Discriminant Analysis (LDA) algorithm computed the optimal weight vector to discriminate between the two classes, which was then used to predict the correct class for every trial in the test set. Decoding performance was quantified as the proportion of correctly classified trials (using the balanced accuracy metric), averaged across all cross-validation iterations within each participant.

Because (2a-b) are two basic-level categories within each superordinate category, and (3a-d) are four exemplar categories within each basic-level category, the decoding outputs were later averaged together across all pairings to compute a single decoding performance trace for each category level, within each subject. To account for the trial number differences between the superordinate and basic-level category decoders, in a control analysis, for each cross-validation fold of the superordinate decoding analysis, we pseudo-randomly split the trials in half and computed the decoding accuracies separately for each half, and then averaged the two accuracy traces. This analysis produced a similar pattern of results as in Fig 3 A-C and G.

For the cross-image-location decoding analysis in Fig. S5, a similar procedure was followed using the Linear Discriminant Analysis (LDA) algorithm, except that trials were sorted by the 4 image locations (top, bottom, left, right) and trials from 3 out of 4 locations were chosen to be the training set, and trials from the left-out location were chosen to be the test set. The training set was subsampled to achieve not only balanced classes, but also balanced exemplar counts. The trials within the training set were then averaged over randomly chosen groups of 4 trials, and the classifier was tested on each trial of the test set (a “leave-one-trial-out” scheme). This was repeated for 300 iterations, for each of the 4 train-test splits. All resulting balanced accuracies were averaged to compute a single decoding performance trace for each category level and image condition, in each subject.

For the target-image position decoding analysis in Fig. S7, a Linear Discriminant Analysis (LDA) classifier was also used. The classifier was configured for pairwise (1 vs. 1) comparisons across all 4 image locations (e.g., top vs. bottom, top vs. right, top vs. left, left vs. bottom, etc.) and all resulting balanced accuracies were averaged for each image condition.

### Time Generalization-Matrix

To examine the temporal stability of the superordinate category representations, a time generalization decoding analysis was performed (Fig. S6) (King and Dehaene, 2014). The previously described decoding analysis was repeated, except that the classifier was trained at each timepoint and tested on all timepoints in the trial epoch, yielding a two-dimensional train × test time decoding accuracy matrix. Similar to the description above under “Multivariate Pattern Analysis”, a stratified cross-validation procedure was used with balanced exemplar counts in the training set, and repeated for 300 iterations.

### Channel-level searchlight analysis

To shed light on the potential spatial distribution of involved neural activity, an exploratory channel-level searchlight decoding analysis was performed. Within groups of neighboring channels, the previous decoding analyses were repeated to classify the superordinate (animal vs. vehicle) and basic-level categories (dog vs. cat, and car vs. truck). The two basic-level decoding performances were later averaged. For each subject, neighboring channels were selected using the triangulation approach (on average 7.4 channels per neighborhood). Consistent with the procedure described above, for each searchlight neighborhood, a stratified cross-validation procedure with 100 iterations was implemented. The resulting performance was stored at each center channel. We then computed group-level statistics across both space and time using cluster-based permutation testing (see “Decoding analysis statistics” below for details). Across space, searchlight channel neighbors were determined using each channel’s immediately adjacent channels via the triangulation approach. Across time, clusters were determined by groups of consecutive timepoints.

### Decoding analysis statistics

To identify significant time points and/or channel clusters, cluster-based permutation testing was implemented using CoSMoMVPA. At each timepoint, one-sample or paired t-tests were computed against the null hypothesis (chance-level at 50% for binary classification, or 0% for the differences between two decoding accuracy traces). A null distribution was generated using 10,000 Monte Carlo sign-flip permutations, in which the sign of each participant’s accuracy trace was randomly flipped. For each permutation, clusters of contiguous time points and/or neighboring channels were identified (with a cluster-defining threshold of p < 0.05), and the cluster statistic was computed as the maximum cluster size. Observed clusters were marked as significant if their size exceeded the 95^th^ percentile of the null distribution.

### Bootstrapping statistics for onset and peak decoding times

To compare onset and peak decoding times across category levels (Sassenhagen and Draschkow, 2019), onset and peak times were estimated using a bootstrapping procedure across 20,000 iterations (Cichy et al., 2014; Pernet et al., 2015). For each iteration, the data were resampled with replacement across the 30 participants, and cluster-based permutation testing with 10,000 sign-flip permutations was performed, from which the first significant timepoint (onset) and the timepoint at maximum decoding accuracy (peak) were extracted. 95% confidence intervals were computed on the resulting onset and peak latency distributions. Overlapping confidence intervals were interpreted as indicating non-significant differences in onset or peak decoding times between category levels.

### Bayesian Statistics for Decoding Analysis

Bayesian hypothesis tests were performed on select analyses yielding null results to quantify the evidence in favor of chance-level decoding performance. At each time point, participant-level decoding accuracies were compared to the chance-level performance (0.5) using a one-sample Bayesian t-test, implemented in the “BayesFactor” package from R (Morey, 2018). Following recommendations from Teichmann et. al. (2022) (Teichmann L., 2022), a half-Cauchy prior (with scale parameter r = 0.707) was used. A null interval was defined as δ ≤ 0.5, and effect sizes below this threshold were treated as practically equivalent to chance. Bayes Factors (BF) were interpreted as substantial evidence for above-chance performance when log_10_(BF) > 0.48 (BF > 3), and for chance-level performance when log_10_(BF) < -0.48 (BF < 1/3).

### Neural Geometry Analysis: cross-category generalization performance (CCGP) and parallelism score (PS)

To test for the abstraction of the superordinate category representation, disentangled from low-level object features, a cross-category decoding analysis was performed using Linear Discriminant Analysis (LDA), separately for each image condition (Normal, Masked, and LSF). Here, basic categories were arranged across the animacy boundary (cat vs. truck, cat vs. car, dog vs. truck, dog vs. car), and classifiers were trained on one category pair and tested on a different pair (e.g., trained on cat vs. truck and tested on dog vs. car, see Fig 4A). As a control, classifiers were trained and tested within the same category pair.

Decoding was carried out using the stratified cross-validation procedure for 100 iterations, as described above in “Multivariate Pattern Analysis.” Within each iteration, trials were first balanced within each participant’s dataset by randomly subsampling to match the minimum class count at the exemplar level, identical to the procedure described above. For the cross-category generalization performance (CCGP) (Bernardi et al., 2020) analysis, training and testing partitions were determined by the pairwise animacy groupings, as described above and illustrated in Fig. 4A. Then, CCGP was computed by averaging decoding accuracies across all four possible train-test pair combinations. For the control analysis, decoding performance was evaluated by training and testing within the same pairwise animacy groupings (e.g., trained and tested on cat vs. truck) using a stratified cross-validation approach. Here, 90% training and 10% testing partitions were created by assigning approximately 10% of the total trials to the test set in multiples of the full exemplar set (i.e., in multiples of 6 exemplars), ensuring that the training set retained balanced exemplar counts. Finally, for both CCGP and control analyses, decoding performance was quantified as the proportion of correctly classified trials (using the balanced accuracy metric), averaged across all cross-validation iterations in each participant.

For the cross-image location CCGP decoding analysis, a similar procedure was followed using the Linear Discriminant Analysis (LDA) algorithm with a leave-one-trial-out scheme. Trials were sorted by the 4 image locations (top, bottom, left, right) and trials from 3 out of 4 locations were chosen to be the training set, and trials from the left-out location were chosen to be the test set. The training set was balanced by sub-sampling to achieve equal class and exemplar counts and the classifier was tested on each trial of the test set. This procedure was iterated 100 times for each of the four train-test splits. All resulting balanced accuracies were averaged to compute a single decoding performance trace for each analysis (CCGP and control, for each image condition), in each subject.

For each image condition’s CCGP analysis, the degree of abstraction was also examined via the Parallelism Score (PS), derived from the classifier weight vectors. For each participant and time point, weight vectors corresponding to each basic category pair were extracted. Weight vectors are orthogonal to the hyperplane in the linear classifier that maximally separates the two classes. The similarity between pairs of weight vectors (e.g., between dog vs. truck and cat vs. car) reflects the alignment of their representational geometries—the more parallel, the more aligned (Bernardi et al., 2020; Ito et al., 2022; Courellis et al., 2024) (see Fig. 4E for an illustration schematic).

Weight vectors were normalized to capture the direction, regardless of the magnitude, and cosine similarities between pairwise weight vectors, w_i_ and w_j_, were computed as cos(*θ_i_*_,*j*_) = w*_i_* · w*_j_*. The parallelism score (PS) was computed as the mean cosine similarity across 2 pairwise weight vector combinations (comparing the weight vectors corresponding to the red lines in Fig. 4A, the following cosine similarities are computed: (1) between cat/car and dog/truck, and (2) between dog/car and cat/truck). Values closer to 1 indicate more parallel (aligned) hyperplanes, and values closer to 0 indicate orthogonal hyperplanes. PS time series were baseline-corrected by subtracting the mean value over the pre-stimulus interval (-0.4 to 0 s). Higher PS values indicate a neural representational structure that supports a more abstract superordinate category representation (e.g., the representation of “animal” regardless of whether it’s a dog or a cat).

Statistically significant timepoints of the Parallelism Score, where hyperplanes were considered significantly aligned, were assessed using cluster-based permutation testing in CoSMoMVPA. At each timepoint, one-sample t-tests were computed against 0 (the expected value for orthogonal weight vectors). A null distribution was generated using 10,000 Monte Carlo sign-flip permutations, in which the sign of each subject’s PS trace was randomly flipped. For each permutation, clusters of contiguous time points were identified (with a cluster-defining threshold of p < 0.05), and the cluster statistic was defined as the maximum cluster size. Observed clusters exceeding the 95^th^ percentile of the null distribution were considered statistically significant (p < 0.05, one-tailed).

A Bayesian hypothesis test was also performed at each time point to quantify the evidence for the null hypothesis (PS = 0), using parameters described above in “Bayesian Statistics for Decoding Analysis.”

### Distance-to-bound analysis

To assess how representational strength for each category level contributes to each trial’s reported awareness level, a distance-to-bound analysis was implemented. EEG channel voltages from all channels were used to decode superordinate (animate vs. inanimate) and basic categories (dog vs. cat or car vs. truck, separately within each superordinate category) using linear discriminant analysis (LDA) classifiers. The analysis followed the same procedure as the decoding analysis described above, but employed a leave-one-trial-out cross-validation scheme and the inclusion of all image conditions into the classifier. To balance exemplar counts within each training class, trials were subsampled to match the minimum exemplar count. Here, the procedure balanced 12 exemplars across all 3 image conditions for a total of 36 exemplar types for the superordinate classifier, or 6 exemplars across all 3 image conditions for a total of 18 exemplar types for the basic-level classifier. This process was repeated 100 times for every left-out trial. Distances corresponding to the same test trial were averaged across iterations.

In each iteration, the distances were calculated by projecting each test trial’s feature vector x (i.e., preprocessed EEG data across all channels) at each time point onto the discriminant dimension defined by the weight vector w, with offset w_0_, as follows:

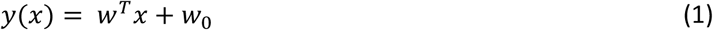

The resulting value, *y*(*x*), represents the distance of each trial from the decision boundary in the projection that maximally separates the two classes. Distance values across time are visualized as the green double-sided arrows and trace in Fig. 5A.

To interpret superordinate or basic level distances across the two discriminated classes, the values were converted to “signed distances” by multiplying the distances associated with the 2^nd^ class (as determined by the “ground-truth” label) by -1, following previous studies (Ritchie et al., 2015; Contini et al., 2021). In Eq (1), a positive *y*(*x*) value would indicate that the classifier chooses class 1, whereas a negative *y*(*x*) value would indicate that the classifier chooses class 2. Thus, the above procedure (multiplying *y*(*x*) by - 1 whenever the true label is the 2^nd^ class) ensures that correct classifier choices are always associated with a positive signed distance, whereas incorrect classifier outputs are always associated with a negative signed distance. In addition, the magnitude of the signed distance reflects the strength of neural evidence for the classifier choice. In using the signed distances, we specifically assess category representations that are congruent with sensory evidence.

To test how superordinate or basic-level signed distances predicted PAS ratings, we fit a cumulative-link mixed model (CLMM) at each timepoints (Visalli et al., 2024). Separately for signed distances derived from the superordinate and basic level classifiers, the model was fit using 30 ms sliding window in steps of 10 ms, using R’s “ordinal” package (Christensen, 2023). Signed distance values were first z-scored within each subject. With Y as the trial-level PAS rating for each subject *i* and for each trial *k,* the relationship between PAS ratings and signed distances is modeled for each timepoint. This model was fit separately for the superordinate and basic level classifiers, as follows:

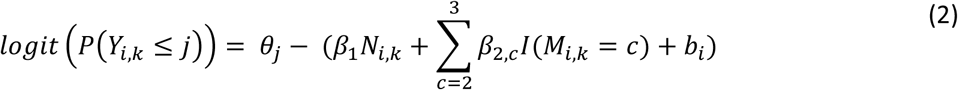

where ordinal PAS ratings Y were modeled as a latent continuous variable, ranging from no subjective experience (PAS = 0) to a vivid subjective experience (PAS = 3). Threshold parameters *θ*_j_ were estimated, where j indexes the cumulative logit thresholds, such that the thresholds divide the latent variable into the 4-point PAS rating scale (i.e., corresponding to be boundaries between 0/1, 1/2, and 2/3). Proportional odds are assumed (i.e., that predictor effects are constant across PAS rating thresholds). The model included two fixed effects: for each subject *i* and for each trial k*, N_i_*_,*k*_ denotes the signed distance from the classifier output (computed separately for the superordinate and basic-level classifiers). β_1_ is the fixed effect coefficient for *N_i_*_,*k*_, which quantifies how signed distances influence the latent PAS rating, and is plotted across time in Fig. 5E. *M_i_*_,*k*_ denotes image conditions (Normal, Masked, and LSF) as a factorized variable, where *I*(*M_i_*_,*k*_ = *c*) is 1 if trial k is image condition c, and 0 otherwise; c = 2 or 3 corresponds to the image conditions (Masked and LSF) compared to the Normal condition as the reference level. *β*_2,*c*_ consists of two fixed effect coefficients, *β*_2,2_ and *β*_2,3_ for the two image condition comparisons, respectively (Masked vs. Normal for c = 2; LSF vs. Normal for c = 3). These terms were included as a nuisance regressors to control for image-condition specific effects. *b_i_* is a subject-specific random intercept.

For this analysis, trials with no PAS response where the participant skipped the trial were excluded. In addition, only trials with correct superordinate categorization answers are included in this analysis. This analysis choice strikes a balance between removing potential trials with lapses of attention and keeping a wide range of PAS ratings. Nevertheless, control analyses using all trials or using only trials with correct basic-level categorization performance yielded broadly consistent results.

Group-level significance was assessed using a cluster-based permutation test applied to the model coefficient β_1_ (corresponding to the signed distance). At each 30 ms time window, a Wald z-statistic (the coefficient divided by its standard error) for β_1_ was computed. Temporal clusters were defined as contiguous time windows where |z| > 1.96 (p < 0.05 uncorrected). The cluster-level statistics were computed as the number of contiguous significant time bins. To generate a null distribution of cluster sizes, the signed distances were randomly shuffled across trials within each participant 1000 times, preserving the temporal structure within each trial and the relationship between the PAS rating and image condition. The model was fit for each permutation as before, and empirically observed clusters exceeding the 95^th^ percentile of the null distribution were considered statistically significant (p < 0.05).

To test how superordinate or basic-level signed distances predicted the categorization behavior, we follow a similar approach as above, except that we fit a logistic mixed-effects model for each category level to predict the binary correct/incorrect outcomes at each timepoint. To control for subjective awareness, only PAS=1 trials (which had the highest trial count, see Fig. S2G) were used in this analysis.

The equation is as follows:

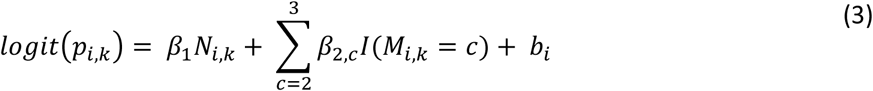

Here, *p_i_*_,*k*_ is the superordinate or basic-level categorization response (0 for incorrect; 1 for correct) for every subject i and for every trial k; *N_i_*_,*k*_ denotes the signed distance from the classifier output (computed separately for the superordinate and basic-level classifiers). β_1_ is the fixed effect coefficient for *N_i_*_,*k*_, which quantifies how much the signed distances influence categorization performance (superordinate signed distances are used to predict the superordinate categorization performance; basic-level signed distances are used to predict the basic-level categorization performance). *M_i_*_,*k*_ denotes image condition (Normal, Masked, and LSF) as a factorized variable, where *I*(*M_i_*_,*k*_ = *c*) is 1 if trial k is image condition c, and 0 otherwise; c = 2 or 3 corresponds to the image conditions (Masked and LSF) compared to the Normal condition as the reference level. *β*_2,*c*_ consists of two fixed effect coefficients, *β*_2,2_ and *β*_2,3_ for the two image condition comparisons, respectively (Masked vs. Normal for c = 2; LSF vs. Normal for c = 3). *b_i_* is the subject-specific random intercept.

Group-level significance was assessed using a cluster-based permutation test applied to the model coefficient β_1_ (corresponding to the signed distance), similar to the previous paragraph describing the approach for the cumulative-linked mixed effects model.

### Mediation analysis

As a follow-up to the distance-to-bound analysis, mediation analyses were conducted to assess whether superordinate or basic level signed distances independently influenced PAS ratings (X → Y), or whether one variable mediated the effect of the other (X → M → Y). Trial-wise signed distance values were extracted at the peak timepoint of each of the main significant clusters from CLMM results in Fig. 5E: early basic (at 62 ms), early superordinate (at 36 ms), late basic (at 368 ms), and late superordinate (at 363 ms). The peak signed distance values were z-scored across trials within each participant. The peak distance of the late superordinate cluster was assigned as the mediating variable M, and the other peak distances were assigned as the independent variables X in the three mediation models, respectively, as described in Table 2.

**Table 2:** Mediation analysis design (corresponding to. **Fig. 6).**

|  | Independent Variable X | Mediating Variable M | Outcome Variable Y |
| --- | --- | --- | --- |
| **Model 1 (Fig. 6B)** | Early Basic (62 ms) | Late Superordinate (363 ms) | PAS rating |
| **Model 2 (Fig. 6C)** | Late Basic (368 ms) | Late Superordinate (363 ms) | PAS rating |
| **Model 3 (Fig. 6D)** | Early Superordinate (36 ms) | Late Superordinate (363 ms) | PAS rating |

For each model, the total effect of X on Y was decomposed into direct and indirect pathways, as shown in Fig. 6A. Path **a** represents the effect of the independent variable X on the mediator M, estimated using linear regression, separately for each participant, as follows:

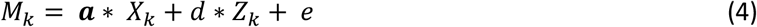

*M_k_* is the peak signed distance of the mediator variable M for each trial *k, X_k_* is the peak signed distance predictor variable of the independent variable X for each trial *k*, and **a** is the fixed effect coefficient for *X_k_.Z_k_* is the factorized conditions variable (Normal, Masked, and LSF) for each trial *k*, and d is the fixed effect coefficient for *Z_k_*. d was considered a nuisance regressor to control for the confounding effects of the image condition on the relationship between X and M. *e* represents the residual error.

Path **b** represents the effect of the mediator M on the outcome Y, controlling for X, and path **c’** represents the effect of the independent variable X on the outcome Y, controlling for M. Paths **b** and **c’** were modeled using a cumulative linked model (CLM), fit separately for each participant, as follows:

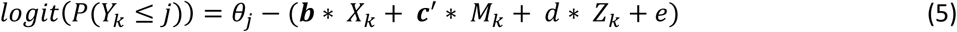

Similar to equation 2, CLM models the ordinal outcome Y, the PAS ratings, as a latent continuous measure, and the model estimates threshold parameters *θ*_j_ that divides the latent measure into a 4-point PAS rating scale. *M_k_* is the peak signed distance of the mediator M or each trial *k,* and **b** is the fixed effect coefficient for *M_k_*. *X_k_* is the peak signed distance predictor variable of the independent variable X for each trial *k,* and **c’** is the fixed effect coefficient for *X_k_*. *Z_k_* is the factorized conditions variable (Normal, Masked, and LSF) for each trial *k*, and d is the fixed effect coefficient for *Z_k_*. d was considered a nuisance regressor to control for confounding effects of the image condition on PAS ratings. *e* represents the residual error.

For each participant, regression coefficients for paths **a**, **b**, and **c’** were estimated independently. The indirect effects were estimated using the product of coefficients approach by computing **a*b** (Liu et al., 2015). Total effects (c) were computed as the sum of direct (**c’**) and indirect (**a*b**) effects. At the group level, coefficients were tested against 0 using a Wilcoxon signed-rank test (Baron, 1986; Iemi et al., 2022; Koenig and He, 2025). Significant indirect effects (a*b ≠ 0, p < 0.05) indicated mediation.

## RESULTS

### Task Paradigm

We recorded high-density EEG (120-channel) from thirty human participants while they performed a threshold-level object recognition task. On each trial, an image containing an object on a naturalistic background was presented for 60 ms at one of four randomly chosen locations. Phase-scrambled images were displayed in the other three locations. The images were presented in one of three conditions: 1) ‘normal’ condition, where an unaltered original image is presented; 2) visual masking, to selectively disrupt recurrent processing; and 3) LSF (low spatial frequency) condition, in which a low-pass spatially filtered version of the image is presented to degrade feedforward processing (**Fig. 1A**). All target images, regardless of image condition, were presented at threshold contrast (see below), and appeared with four surrounding dots at each corner. Normal and LSF images were followed by a 300 ms blank screen, and Masked images were followed by a 300 ms presentation of the four dots, rotated 90°, as a temporally trailing visual mask. Following extensive piloting (for details, see **Supplementary Results**), visual masking in this experiment was achieved using an Object Substitution Orientation-Shifted mask (OSOM). Similar to the standard object substitution mask (OSM) (Goodhew et al., 2013), OSOM is characterized by non-overlapping target and mask stimuli, which keeps the feedforward sweep as intact as possible while still disrupting recurrent processing.

The stimuli set consisted of 12 exemplar images belonging to 4 basic categories (Dog, Cat, Car, Truck) and 2 superordinate categories (animal/animate vs. vehicle/inanimate) (**Fig. 1B**). The exemplar images were chosen such that the object position, pose, and orientation were well-matched between the two basic categories within each superordinate category (i.e., between cat and dog, and between car and truck). In addition, the images were processed using the SHINE toolbox (Willenbockel et al., 2010) to equate low-level features, including luminance and spatial frequency, across all original exemplar images.

**Figure 1:**
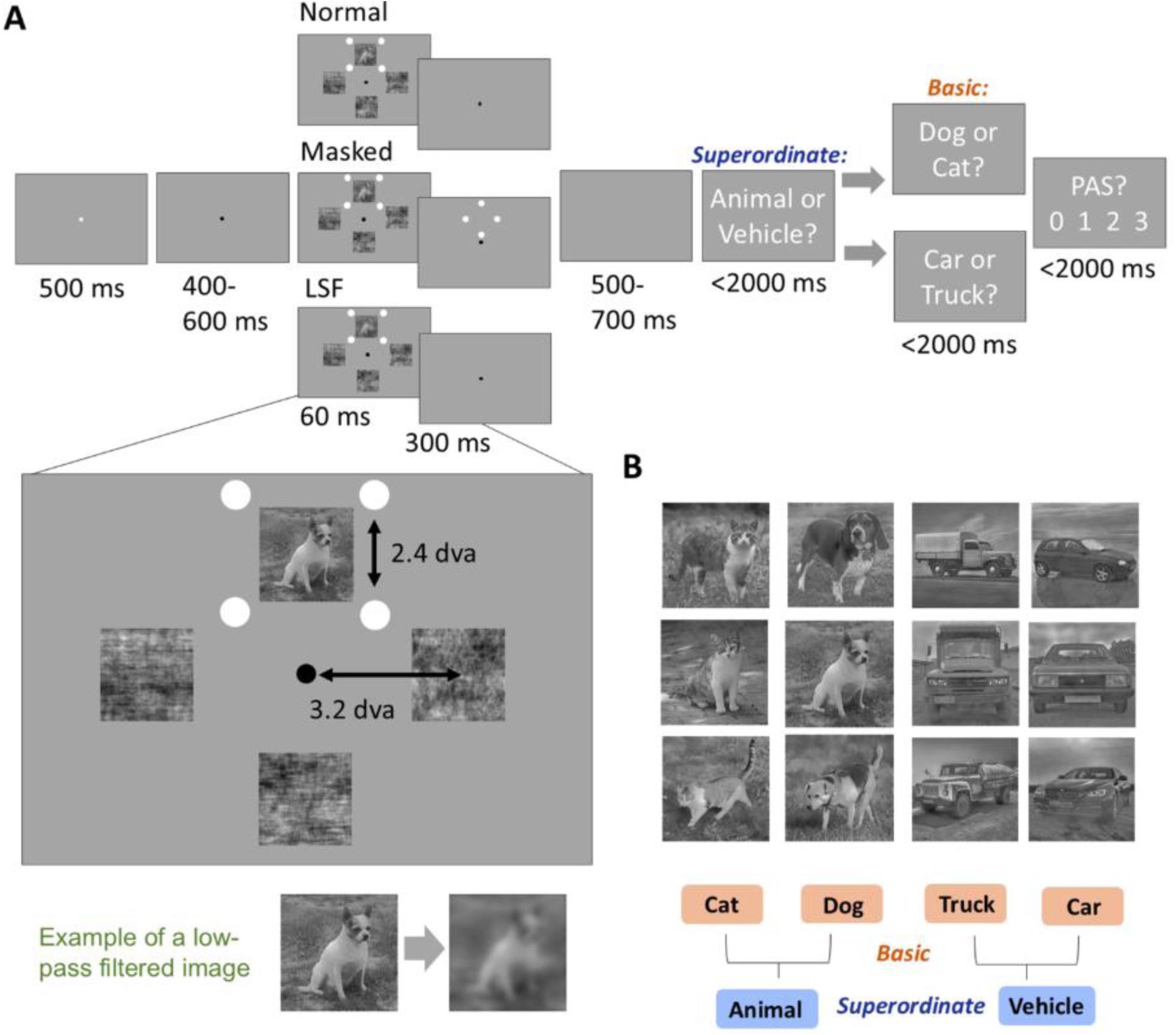
Paradigm and hypothesis. **(A)** Top: Trial structure of the main object categorization task. Images were presented for 60 ms in one of three image conditions (Normal, Masked, or LSF), followed by two categorization questions and a subjective vividness report using the perceptual awareness scale (PAS) ratings from 0 (“no experience”) to 3 (“clear experience”). The target image location was randomly chosen between top, bottom, left, and right locations with equal probability, and filler images (phase-scrambled images) were placed in the remaining locations. Bottom: An example of a low spatial frequency (LSF) image. **(B)** 12 exemplar images used in the task, which belong to 4 basic-level categories (cat, dog, truck, and car) and 2 superordinate categories (animal and vehicle). Image luminance and spatial frequency content were matched across stimuli, and the contrast of each image was titrated for each participant using a staircasing procedure based on the individual performance for the basic-level categorization task in the normal condition.

On each trial, the specific exemplar image, image location, and condition were pseudo-randomly chosen with equal probability. Prior to the main task, the contrast of each original exemplar image was titrated for each participant to reach the individual threshold, defined as ∼63% basic-level categorization performance in the Normal condition, using a staircasing procedure (see Methods for details). The same image contrast was also used in the LSF and Masked conditions, and the phase-scrambled counterparts were presented at the same titrated contrasts.

Following each image presentation, participants answered two 2-alternative forced-choice (2AFC) questions regarding the object category: first the superordinate category (“Animal or Vehicle?”) and then the basic category (“Dog or Cat?” if they answered “Animal”; “Car or Truck?” if they answered “Vehicle”) (**Fig. 1A**). If the participant responded with the incorrect superordinate category, invalid basic category options were displayed to be congruent with their initial answer. In all following analyses, basic-level categorization performance is computed only on correct superordinate categorization trials, to assess performance only on trials with valid basic-category choices. This analysis choice creates a symmetric 2AFC paradigm format across both category levels, with 50% as the chance-level performance.

At the end of each trial, participants were asked to report the vividness of their perceptual experience via the Perceptual Awareness Scale (PAS) (Overgaard and Sandberg, 2021), a well-validated approach to measure subjective perceptual awareness (Sandberg et al., 2010). A PAS rating of 0 corresponded with “I did not see the target image on the screen, not even the faintest sensation”, a rating of 1 corresponded with “I caught a glimpse of ‘something there’ but I’m not sure what I saw”, a rating of 2 corresponded with “I got a sense of what the presented stimulus might be”, and a rating of 3 corresponded with “I saw the stimulus clearly, without any doubt.”

### Superordinate categorization behavior is relatively robust to the degradation of perceptual awareness

We first examined categorization task behavior. Both superordinate and basic categorization performances were significantly above chance in all three conditions (all p < 0.0001, Wilcoxon signed-rank test, FDR-corrected; **Fig. 2A-B**). A two-way [image condition (Normal, Masked, and LSF) × category level (Superordinate and Basic)] repeated-measures ANOVA on categorization performance revealed significant main effects of both the image condition (F_1,29_ = 169.9, p = 5.65e-25, η_p_^2^ = 0.85) and category level (F_1,29_ = 516.4, p = 5.01e-20, η_p_^2^ = 0.95), as well as a significant interaction effect (F_1,29_ = 24.7, p = 1.73e-8, η_p_^2^ = 0.46). These results show that the superordinate categorization performance is more robust than the basic-level categorization performance, and that image manipulations had a greater effect on basic-level categorization.

**Figure 2:**
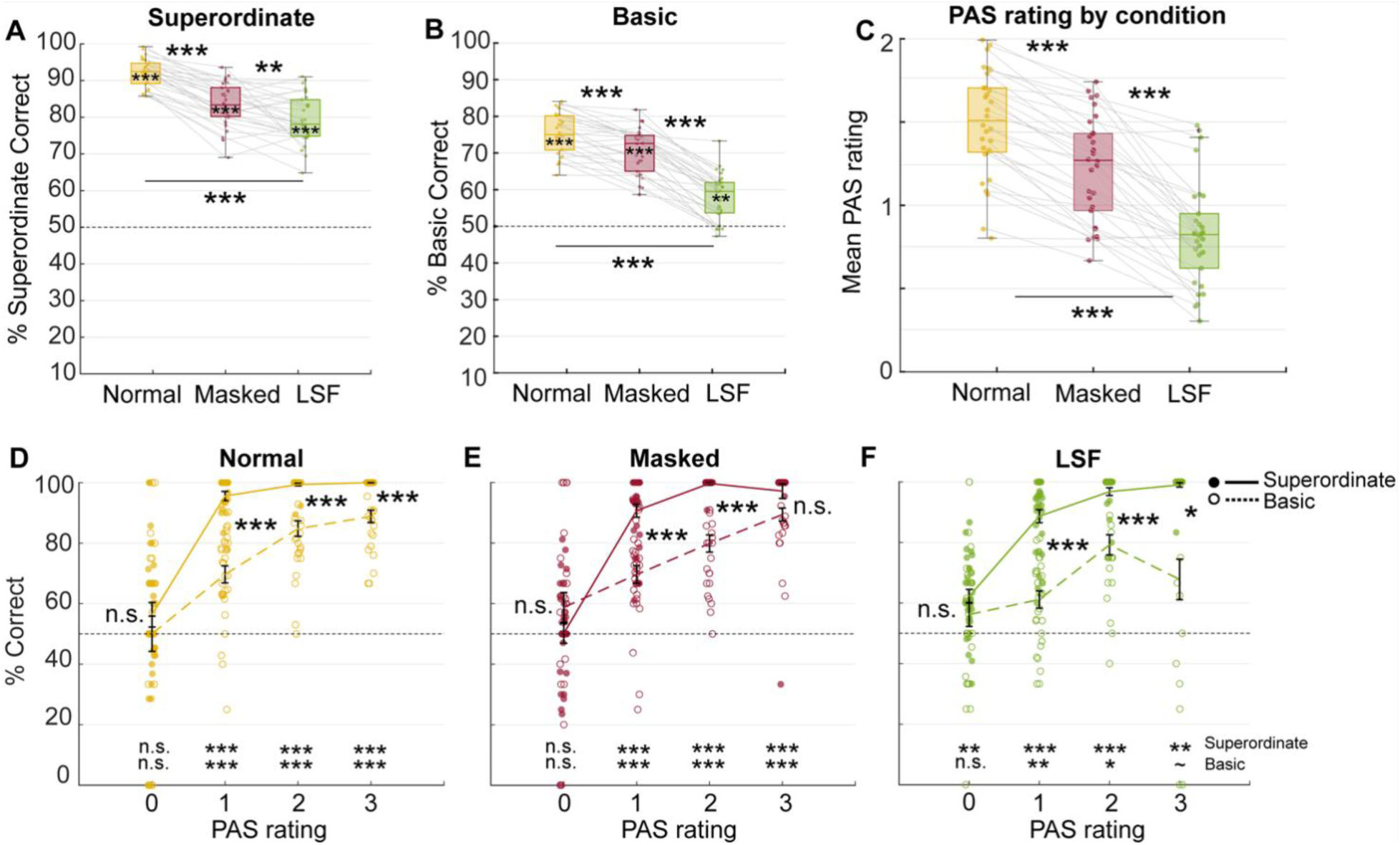
Behavior results from the Main Experiment with EEG. **(A)** Categorization performance results across three image conditions for superordinate category responses. The box plots show the data median (middle line) as well as the lower and upper quartiles (top and bottom of the box). Each dot represents an individual subject’s averaged performance across trials, per condition, and the light gray lines connect categorization performances across conditions within subjects (n = 30). **(B)** Same as (A) but for basic category responses. Only correct superordinate trials (which prompted valid basic category questions) are included. **(C)** Mean PAS ratings across three image conditions (n = 30). All asterisks between box plots indicate significance based on Bonferroni-corrected post-hoc statistics. Asterisks printed on top of the box plots are statistics from Wilcoxon signed-rank tests against 50% chance-level performance (FDR-corrected). * p < 0.05, ** p < 0.01, *** p < 0.001. **(D-F)** Superordinate (solid line) and basic (dashed line) categorization performances, split by PAS ratings, for each of the image conditions. Each dot represents an individual subject. Asterisks indicate Bonferroni-corrected post-hoc comparisons between the superordinate and basic-level categorization performances, for each image condition and PAS rating. For all plots, the black dashed line indicates chance-level accuracy (50%). Asterisks printed at the bottom indicate statistics from Wilcoxon signed-rank tests against 50% chance-level performance, FDR-corrected. ∼ p < 0.08 (marginal effects), * p < 0.05, ** p < 0.01, *** p < 0.001.

We next tested whether categorization performance varied by stimulus category. A two-way [image condition (Normal, Masked, and LSF) × superordinate category (animal vs. vehicle)] repeated-measures ANOVA revealed a significant effect of category (F_1,29_ = 6.0, p = 0.021, η_p_^2^ = 0.171; **Fig. S2A**). Post-hoc tests using Bonferroni correction revealed that this was driven by the masked condition where vehicles had significantly higher categorization performance than animals (p = 0.001). However, another two-way (image condition × basic-level category) repeated-measures ANOVA revealed a non-significant effect of basic-level category (F_1,29_ = 2.03, p = 0.116, η_p_^2^ = 0.065, **Fig. S2B**). Together, these results show that categorization performance was largely comparable across stimulus categories.

We then examined image manipulations’ effects on perceptual awareness. A one-way repeated-measures ANOVA on PAS ratings revealed a significant main effect of condition (F_1,29_ = 96.22, p = 3.78e-19, η_p_^2^ = 0.77; **Fig. 2C**). Post-hoc tests using the Bonferroni correction revealed significant differences between the Normal and Masked conditions (p = 2.11e-8), the Masked and LSF conditions (p = 6.02e-9), and the Normal and LSF conditions (p = 1.35e-19). The overall trend is similar to categorization performance: the Normal condition had the highest PAS rating, followed by the Masked condition, and then the LSF condition.

We then tested whether stimulus category influenced PAS ratings using two-way (image condition × superordinate/basic category) repeated-measure ANOVAs. In both ANOVAs, the effect of category was significant (superordinate: F_1,29_ = 10.86, p = 0.003, η_p_^2^ = 0.272; basic: F_1,29_ = 10.86, p = 0.003, η_p_^2^ = 0.272). Post-hoc tests revealed that this was driven by vehicles triggering slightly higher PAS ratings than animals in the Normal and Masked conditions (**Fig. S2C**), which may be due to the relative salience of straight edges and lines in vehicles (which are not removed by the SHINE algorithm applied to equalize luminance and spatial frequency). However, post-hoc tests revealed non-significant pairwise comparisons between basic-level categories in 17 out of 18 pairwise comparisons (**Fig. S2D**).

Finally, we examined how categorization performance varies with viewing condition and perceptual awareness by conducting a 3-way repeated-measures ANOVA on categorization performance [factors: image condition (Normal, Masked, LSF) × category level (Superordinate and Basic) × PAS rating (0 to 3)] (**Fig. 2D-F**). This analysis revealed significant main effects for category level (F_1,1_ = 57.9, p = 6.2e-6, η_p_^2^ = 0.77) and PAS rating (F_1,3_ = 37.8, p = 3.4e-8, η_p_^2^ = 0.67), and a significant category level × PAS rating interaction effect (F_1,3_ = 8.2, p = 0.001, η_p_^2^ = 0.77). No other main or interaction effects were significant, including the main effect of condition (F_1,2_ = 1.5, p = 0.24, η_p_^2^ = 0.77) and the image condition x category level interaction (F_1,2_ = 2.2, p = 0.14, η_p_^2^ = 0.97).

Notably, the superordinate categorization performance reaches near-ceiling level at PAS ≥ 1 and is significantly higher than the basic-level categorization performance starting at PAS ≥ 1 (p < 0.001, post-hoc comparisons between category levels at PAS = 1, FDR-corrected; **Fig. 2D-F**). A PAS rating of 1 corresponds to a weakly conscious gist perception — “I only caught a glimpse”; thus, superordinate categorization performance is relatively robust at low awareness levels.

Strikingly, even in unaware trials (with PAS ratings of 0, indicating no conscious detection of the target image), superordinate categorization performance in the LSF condition was significantly above the chance level (62%, p = 1.13e-4, Wilcoxon signed-rank test against 50% chance, FDR-corrected). All other categorization performances were at chance level at PAS = 0 (**Fig. S2E-F**). This result suggests that superordinate categorization performance, when driven by low-frequency spatial information alone, does not require conscious awareness.

Together, these behavioral findings establish that superordinate categorization performance is relatively robust to image degradation and poor awareness, whereas basic-level category representations emerge with more vivid experiences.

### Early onset of both superordinate and basic-level category information

To elucidate the sequence of neural events underlying conscious object perception at the category level, we first performed a multivariate decoding analysis using EEG signals at each time point to decode superordinate (animal vs. vehicle) and basic (dog vs. cat; car vs. truck) object categories in each condition separately, pooling trials across all image locations (top, bottom, left, right) together. Significant superordinate and basic category representations emerged at similar times across all conditions, where onset times showed overlapping confidence intervals: 185 ms [95% confidence interval: 90–320 ms], 150 ms [110–505 ms], and 100 ms [-30 – 200 ms] for superordinate category decoding in the normal, masked and LSF condition, respectively (**Fig. 3B**, blue). Onset timing for significant basic-level category decoding was 110ms [85–185ms], 90 ms [70–595ms], and 85 ms [80–595ms] for Normal, Masked, and LSF conditions, respectively (**Fig. 3B**, orange).

**Figure 3:**
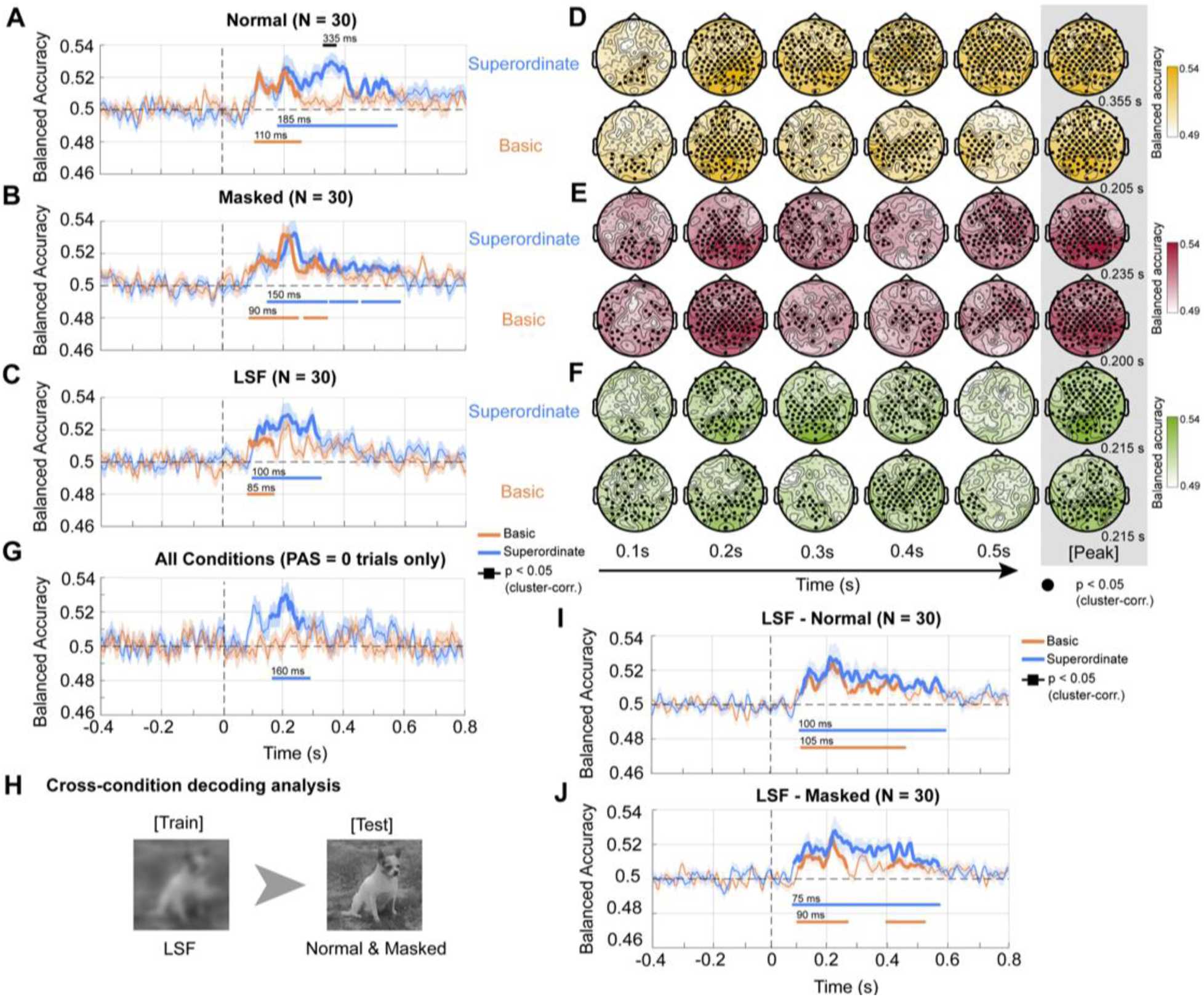
Superordinate and basic category decoding analysis. **(A-C)** Decoding analysis results for superordinate (cat- and-dog vs. car-and-truck) and basic-level (average of cat vs dog and car vs. truck) categories in the Normal, Masked, and LSF conditions. The two decoding accuracy traces for the basic category pairs (cat vs. dog, and car vs. truck) were averaged within each subject. Shaded areas indicate s.e.m. across subjects (*n* = 30). Bolded traces and horizontal lines indicate significant timepoints (p < 0.05, cluster-based permutation test). The black horizontal line above the traces indicates a significant difference between the superordinate and basic category decoding accuracies. Numbers indicate the onset times of significant decoding. **(D-F)** Searchlight decoding analysis at the channel level from t = 0.1 s (100 ms after stimulus onset) to t = 0.5 s, and at peak decoding times (rightmost column) for **(D)** Normal, **(E)** Masked, and **(F)** LSF conditions. Black dots indicate significant channel clusters (p < 0.05, cluster-corrected across space and time). **(G)** Decoding analysis results for superordinate and basic categories on trials with PAS ratings reported as “0” (no perceptual experience), pooling across the 3 conditions. Corresponding Bayesian statistics are shown in **Fig. S4**. **(H)** Schematic for cross-condition decoding analysis. For both category levels, the classifier was trained on LSF images and tested on Normal or Masked images. **(I-J)** Category decoding analysis results for classifiers trained on LSF images and tested on Normal (**J**) or Masked (**I**) images. Format is the same as *A-C*. In all decoding analyses, chance level is 50%.

Peak decoding times also did not differ significantly between superordinate and basic-level categories (**Fig. S3C**). In addition, both onset and peak times of decoding did not significantly differ between superordinate/basic-level category decoding and exemplar-level decoding within a basic category (**Fig. S3A-C**). Overall, this pattern of roughly simultaneous emergence of object-related information at multiple levels (exemplar, basic, superordinate) is consistent with previous studies (Carlson et al., 2013; Cichy et al., 2014). Importantly, because only the target image location, not the remaining scrambled-image locations, was indicated by the surrounding 4 dots (see Fig. 1), the subject had clear knowledge of the target image location. Thus, behavioral responses and the corresponding neural activity are based on the processing of the target image content. Consistent with this, target image location could be robustly decoded from neural activity in all three conditions (**Fig. S7**).

Interestingly, the onset time of both superordinate and basic-level category information was earliest in the LSF condition (at 100 ms and 85 ms, respectively), although not significantly earlier than the other conditions. This is notable given that both superordinate and basic-level categorization performances were lowest in the LSF condition (**Fig. 2A-B**). We hypothesized that the low-spatial-frequency information alone might drive the early emergence of category information. To test this hypothesis, we trained classifiers to decode superordinate or basic categories on LSF images and tested them on Normal or Masked conditions (**Fig. 3H**). In both cases, superordinate and basic categories were successfully decoded with an early onset, with slightly earlier onset times than the original decoding analysis (**Fig. 3I-J**, compared to **3A-B**), supporting our hypothesis. Cross-condition decoding was successful for all pairs of conditions (**Fig. S3F-K**), suggesting that all image conditions carry similar category representations.

Strikingly, the superordinate, but not the basic-level, category representation persisted in unaware trials (where the participants reported PAS ratings of 0 for “no experience of the target at all”; **Fig. 3G**). Bayesian analyses supported this finding: Bayes Factors (BF) exceeded 3 for superordinate category decoding, revealing strong evidence against chance-level performance. Conversely, BF fell below 1/3 for basic-level category decoding, revealing strong evidence for chance-level performance (**Fig. S4A-B**). This finding echoes the behavioral observation of above-chance superordinate categorization performance in unaware LSF trials (**Fig. S2E**), and reveals that superordinate category representations can exist without conscious awareness.

In order to shed light on the potential spatial distribution of involved neural activity, we performed an exploratory channel-level searchlight decoding analysis (**Fig. 3D-F**; for details, see Methods) accompanying the individual condition decoding analysis (**Fig. 3A-C**). Intriguingly, for the superordinate category, successful decoding at 200 ms included prominent clusters overlying the frontal lobe in the Normal and LSF conditions. This finding is consistent with an earlier proposal that the dorsal visual stream’s projection to frontal cortex carries fast, coarse information mediated by the magnocellular pathway, which helps to guide the slower, ventral visual stream processing that fills in the details (Bar et al., 2006). However, this early category representation may or may not directly contribute to perceptual awareness — a question we investigate later.

The above decoding analyses pooled trials across the four image locations; as such, it is unclear if the results were contributed by location-dependent or location-invariant neural code. To answer this question, we separated the trials by target image location (top, bottom, left, right; see **Fig. 1**) and repeated the analyses in Fig. 3A-C and Fig. 3G by training on trials from three image locations and testing on the left-out location. Decoding accuracies were averaged across the four train-test splits. This analysis yielded comparable results to those in Fig. 3 (**Fig. S5A-D**). This result suggests that our category-decoding results are primarily contributed by location-independent neural code. Given the size of our stimuli array (**Fig. 1A**) and known receptive field sizes of different brain regions (Freeman and Simoncelli, 2011; Kay et al., 2015; Riley et al., 2017), these findings suggest that the category representations we decode are largely contributed by high-level visual cortex or prefrontal regions that have relatively large receptive fields.

In sum, these results reveal an overlapping rather than a serial emergence of hierarchical object category representations. Successful cross-condition decoding indicates shared category representations despite differences in the quality of feedforward or recurrent processing. Notably, superordinate category representation was highly significant on unaware trials, when participants denied consciously perceiving any object image.

Although here we have focused on the onset timing of significant category decoding, a prominent feature of these results is that significant superordinate category information persisted much longer in time than basic-level category information (**Fig. 3A-C**). A temporal-generalization matrix for the superordinate category decoding analysis showed that the early and late components of the category information are distinct. This is indicated by the significant clusters that emerge only along the diagonal (**Fig. S6**), suggesting constantly evolving dynamics. Thus, we hypothesized that, rather than being static, the superordinate category neural representation might evolve over time and become more abstract (i.e., less entangled with low-level visual features) later in time. We test this idea and probe the evolving geometry of neural representation next.

### An abstract superordinate category representation emerges during late-stage processing

To test the potential emergence of an abstract superordinate category representation, we adopted two previously established metrics: cross-category generalization performance (CCGP) and parallelism score (PS) (Bernardi et al., 2020; Courellis et al., 2024). CCGP quantifies a decoder’s ability to generalize to task conditions not used for training, and is used here to evaluate whether there is superordinate category representation that is abstracted from the encoding of the specific basic-level category or other low-level object visual features (i.e., an abstract representation of “animal” regardless of whether it is a dog or a cat) (**Fig. 4A**). PS provides additional insight into the geometry of neural representation by quantifying the degree to which the coding directions (which are orthogonal to the respective discriminating hyperplanes) of two classifiers are parallel to each other. The two classifiers are typically constructed to decode the same or related variables in two different conditions (e.g., decoding Cat vs. Car, and decoding Dog vs. Truck) (**Fig. 4E**). Both high CCGP and high PS reflect abstract representations that are generalizable across conditions (for details, see Methods).

**Figure 4:**
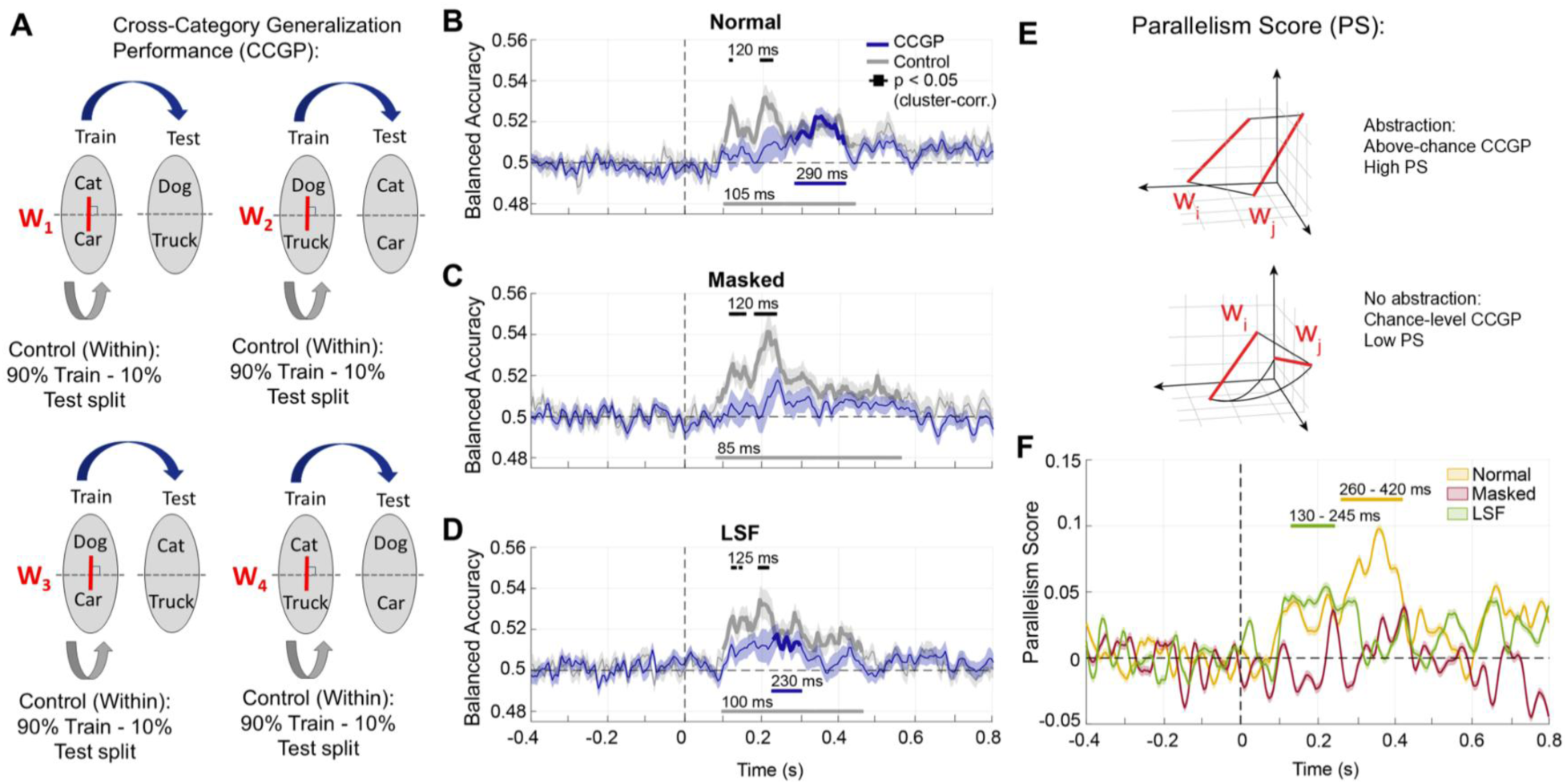
Neural geometry analysis: Cross-category generalization performance (CCGP) and parallelism score (PS). **(A)** Analysis schematic for computing CCGP. Classifiers are trained on one pair of basic categories and tested on a different pair of basic categories, for a total of 4 decoding schemas. For each decoder, the dashed black line denotes the discriminating hyperplane and the red line (**w**) denotes the coding direction. In the control analyses, classifiers are trained and tested within the same basic category pair with a 90%-10% train-test split using a stratified cross-validation procedure (see Methods for details). **(B-D)** CCGP (navy) and control (gray) decoding analysis results for the Normal (**B**), Masked (**C**), and LSF (**D**) conditions. Shaded areas indicate s.e.m. across participants (*n* = 30). The decoding accuracy traces for all pairwise decoding schemas were averaged within each participant. Bolded traces and horizontal lines indicate significant timepoints (p < 0.05; cluster-based permutation test). Black horizontal lines above the traces indicate significant differences between CCGP and control traces. Numbers indicate the onset times of each statistically significant time segment. **(E)** Analysis schematic for computing parallelism scores. The cosine of the angle between two pairwise weight vectors was computed and averaged across relevant pairs of classifiers. Weight vectors specify the coding directions of classifiers and are illustrated by red lines in (**A**) and (**E**). See Methods for details. **(F)** Averaged parallelism scores across participants (n = 30), for Normal (yellow), Masked (red), and LSF (green) conditions. Horizontal lines indicate significant timepoints (p < 0.05; cluster-based permutation test). Corresponding Bayesian statistics are shown in **Fig. S4C-H**.

To calculate CCGP, we trained classifiers to distinguish superordinate categories by training on one basic category pair (e.g., cat vs. truck) and testing whether the classifier generalizes to a different pair (e.g., dog vs. car). We repeated this analysis for all pairwise combinations across the basic categories (**Fig. 4A**). Significant CCGP—indicating abstract superordinate category representations—emerged around 200–300 ms, but only in image conditions with intact recurrent processing: the Normal and LSF conditions (navy traces, **Fig. 4B, D**). This effect was absent in the Masked condition, where recurrent processing was disrupted by the mask (**Fig. 4C**). Bayesian statistical tests confirmed these observations: In Normal and LSF conditions, the Bayes Factor (BF) showed sustained periods with BF greater than 3, providing strong evidence for abstract superordinate category representations. In contrast, the Masked condition mostly showed BF less than 1/3 throughout the time period (**Fig. S4C-E**).

As a control, we trained and tested the classifier on the *same* basic category pairs across the superordinate boundary using a stratified cross-validation approach (i.e., training on a subset of dog vs. truck trials *and* testing on left-out dog vs. truck trials). In this analysis, mid- or low-level features unique to a basic category can drive decoding success (e.g., the cat’s ear). Significant decoding in this control analysis emerged earlier than CCGP (gray traces in **Fig. 4B-D**; **Fig. S4C-E**). In addition, in the early periods (around 100–250 ms), decoding accuracies in the control condition are significantly higher than CCGP decoding accuracies (**Fig. 4B-D**, black bars). These results confirm that abstract superordinate representations emerge later in time than superordinate representations that are entangled with basic-level category representations.

The above analysis pooled together trials with different target image locations. Similar to the decoding analysis, we next repeated the CCGP analysis by training the classifier on trials from three target image locations and testing it on the left-out image location. This analysis yielded broadly similar patterns as the original analysis (compare **Fig. S5E-G** to **Fig. 4B-D**), consistent with our earlier conclusion that the category representation we probe is location-invariant.

To shed light on the spatial distribution of neural activity contributing to the abstract superordinate category representations, we conducted an exploratory searchlight decoding analysis in the channel space (**Fig. S8**). In both the normal and LSF conditions, successful CCGP decoding clusters included substantial frontal electrodes, consistent with known roles of the prefrontal cortex in encoding abstract representations (Bernardi et al., 2020).

Second, to compute the parallelism score (PS), we calculated the cosine of the angle between pairwise weight vectors (Bernardi et al., 2020), with each weight vector representing the coding direction that is orthogonal to the hyperplane separating the two classes for that classifier (**Fig. 4E**; see Methods for details). As with the CCGP analysis, all pairs of basic categories that span the superordinate boundary are evaluated (e.g., decoding cat vs. car, and decoding dog vs. truck; see **Fig. 4A**), with the PS score averaged across them. A higher PS indicates more parallel coding directions, and therefore more aligned representational spaces.

The results reveal significant PS in Normal and LSF conditions, but not in the Masked condition, consistent with the CCGP findings (**Fig. 4F**; Bayesian statistics in **Fig. S4F-H**). Significant PS time segments emerged slightly earlier than the significant CCGP time segments, suggesting that the neural geometry begins realigning along the abstracted superordinate category spaces before a robust generalized representation occurs. Furthermore, significant PS and CCGP scores occurred earlier in the LSF than in the Normal condition, suggesting that low-spatial-frequency information is processed faster.

### Awareness correlates with early basic and late superordinate category representations

Our results thus far reveal superordinate and basic category representations that onset at roughly the same time at ∼100–150 ms, as well as a late abstract superordinate category representation beginning at ∼250–300 ms. In addition, superordinate, but not basic, category representations persist under unaware conditions when participants deny having consciously perceived any object images. We next directly probed the relationship between category representation in the brain and subjective perceptual awareness. To this end, we tested whether the strength of a given category representation at each time point predicts the participant’s reported PAS rating on a trial-to-trial basis. Only trials with correct superordinate categorization were included in this analysis (for details, see Methods).

We operationalized the category-level representational strength as the distance between each trial’s neural activity and the decision boundary in a linear discriminant analysis (LDA) classifier. These distances can be interpreted as the strength of evidence for one class over another. Trials farther from the decision boundary are classified with greater certainty, while those closer to the boundary are less certain. We used a signed distance measure such that correct classifications are associated with positive distances, and incorrect classifications are associated with negative distances (for details, see Methods). A positive relationship between signed distance and PAS rating would suggest that the category representation positively contributes to reported awareness (**Fig. 5A**, see Methods section “Distance-to-bound Analysis” details).

**Figure 5:**
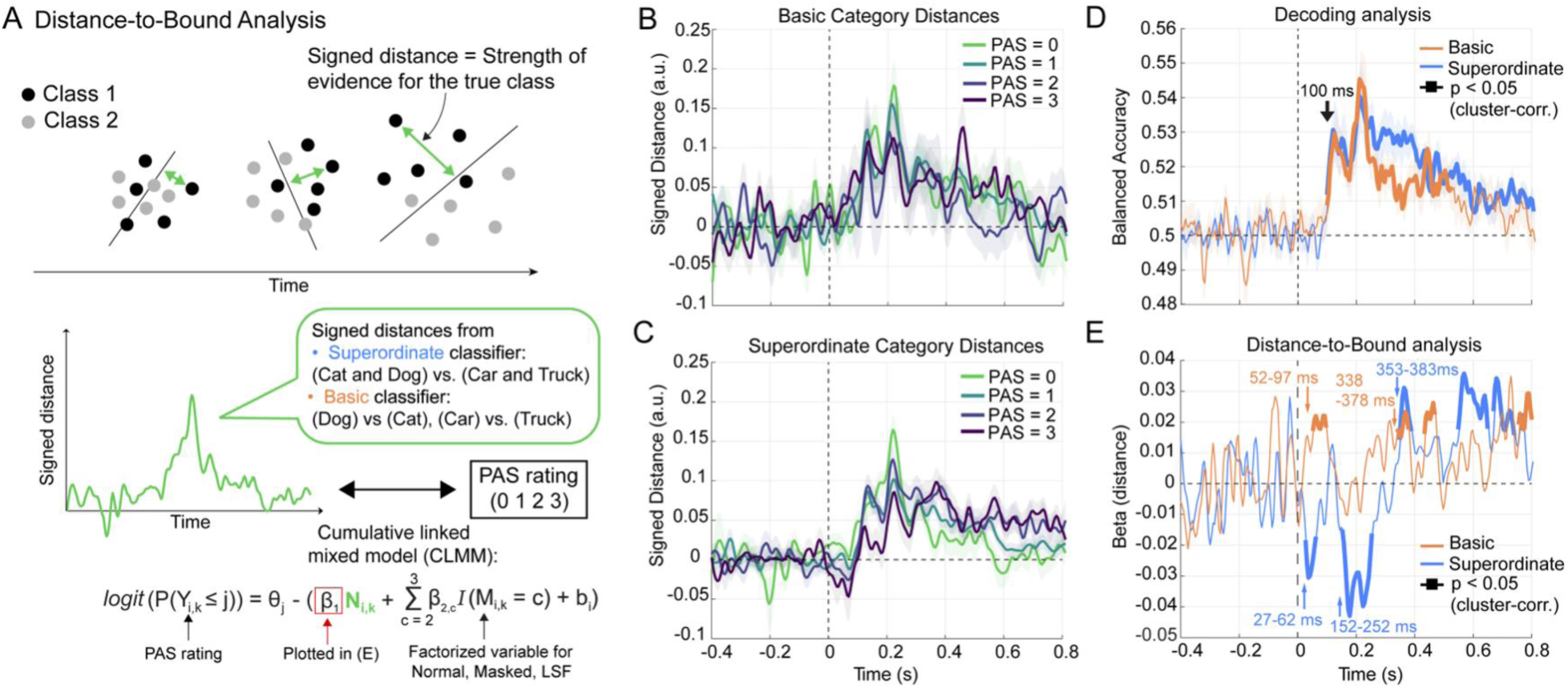
Distance-to-bound analysis testing the contribution of category representations to awareness ratings. **(A)** Analysis schematic for extracting the signed distance of each trial (indicated by the green arrows) from the superordinate (animal vs. vehicle) or basic level (dog vs. cat; car vs. truck) leave-one-trial-out decoding analysis. The corresponding PAS rating was predicted using a cumulative link mixed model. In the equation at the bottom, Y represents the ordinal PAS rating outcome, θ_j_ represents the threshold parameters that partition the PAS ratings, *N_i_*_,*k*_ indicates the signed distance values from the superordinate or basic-level classifier for each subject *i* and each trial *k*, and β_1_ quantifies the effect of signed distance values on PAS ratings and is plotted in (E). *I*(*M_i_*_,*k*_ = *c*) indicates the factorized conditions variable for each image condition (Normal, Masked, and LSF), β_2,2,_ and β_2,3_ are the corresponding coefficients, and b_i_ indicates the subject as a random effect. For details, see Methods. **(B)** Signed distance values for basic-level category, separated by PAS rating. Distance traces were smoothed using a Savitsky-Golay filter with a 150 ms window for visualization. **(C)** Same as (B), but for the superordinate category. **(D)** Decoding accuracy traces for the basic and superordinate level category decoding. Decoding analysis is similar to Fig. 3A-C but was pooled across the 3 conditions. In B-D, shaded area represents the s.e.m. across participants (n = 30). **(E)** β_1_ values for the signed distance variable from cumulative linked mixed model (see Eq. in A), showing the relationship between trial-level signed distances and reported PAS ratings. In D-E, bolded traces indicate significant time points (p < 0.05; cluster-based permutation test). Numbers indicate the onset and offset times of the main significant clusters.

First, in a qualitative analysis, we plot the average time courses for superordinate and basic-level category representation strength (i.e., signed distance to decision boundary), separately for each PAS level (**Fig. 5B-C**). Distinct patterns emerged for the two category levels. For basic-level category representation, trials with higher PAS ratings showed earlier increases in representational strength, such that within 100 ms after stimulus onset, stronger basic-level category representation is correlated with higher PAS ratings (**Fig. 5B**). By contrast, for superordinate category representation, early (0–300 ms) representational distances appeared to be *inversely* correlated with PAS ratings, with higher representational strength for trials reported with low awareness (**Fig. 5C**). After 300 ms, this pattern reverses: trials with high PAS ratings are associated with stronger superordinate category representation.

We used cumulative linked mixed models (CLMMs) to quantitatively assess these patterns (for details, see Methods), with category-level representational strength on individual trials as input data and PAS ratings as the ordinal outcome. The distribution of trial counts for each PAS rating across subjects are shown in Fig. S2G. This analysis confirmed that basic-level category representational strength is significantly positively correlated with PAS rating in an early time period, with a significant cluster at 52–97 ms, followed by a second positive cluster at 338–378 ms (**Fig. 5E**, orange curve). The first cluster immediately precedes or overlaps with the onset of the basic-level category decoding, which reached significance at 100 ms [60 ms – 175 ms] (**Fig. 5D**).

Surprisingly, the strength of superordinate category representation *negatively* correlated with PAS ratings at this early processing stage, with significant clusters at 27–62 ms and 152–252 ms (**Fig. 5E**, blue curve). Similar to the basic-level category representation, these clusters immediately precede or overlap with the onset of the superordinate category decoding, which reached significance at 100 ms [75 ms – 165 ms] (**Fig. 5D**). This result indicates that the weaker the perceptual experiences, the stronger the early superordinate category representation.

This early anti-correlation finding for the superordinate category representation may appear counterintuitive; however, it is consistent with our earlier finding of unconscious superordinate neural representations (**Fig. 3D**) and above-chance superordinate categorization behavior on unaware trials in the LSF condition (**Fig. 2F**; **Fig. S2E**).

Subsequently, for superordinate category representation, a later cluster *positively* correlated with PAS ratings, at 353–383 ms (**Fig. 5E**). This time period overlaps with the emergence of abstract superordinate category representations, as characterized by CCGP and PS metrics in the neural geometry analysis (**Fig. 4**). Thus, the more vivid the perceptual experience, the stronger the late abstract superordinate category representation.

Lastly, we examined the relationship between superordinate/basic-level category representation and the objective categorization accuracy, while controlling for subjective awareness. To this end, we used a logistic mixed-effects model to assess the relationship between neural signed distances and the categorization behavioral performance at a trial-to-trial level (for details, see Methods). To control for subjective awareness, only trials with PAS rating = 1, which had the highest trial count across the 4 PAS rating levels (Fig. S2G), were used in this analysis. This analysis revealed that both superordinate and basic-level neural representational strengths *positively* predicted categorization accuracy in a relatively early time window (47–187 ms for superordinate; 47–97 ms for basic) (**Fig. S9**). Thus, our results reveal an intriguing dissociation between early superordinate category representation’s contributions to objective performance and subjective awareness: positive for the former, and negative for the latter.

### Basic- and superordinate-category representations independently influence perceptual awareness

The above results show that basic-level category representations in both early and late periods, as well as a late abstract superordinate category representation, positively correlate with perceptual awareness. By contrast, an early superordinate category representation negatively correlates with perceptual awareness. To better understand the relationship between these findings, we conducted mediation analyses using trial-level category representational strength as input data and PAS ratings as the predicted ordinal outcome (**Fig. 6A**; for details, see Methods).

**Figure 6.**
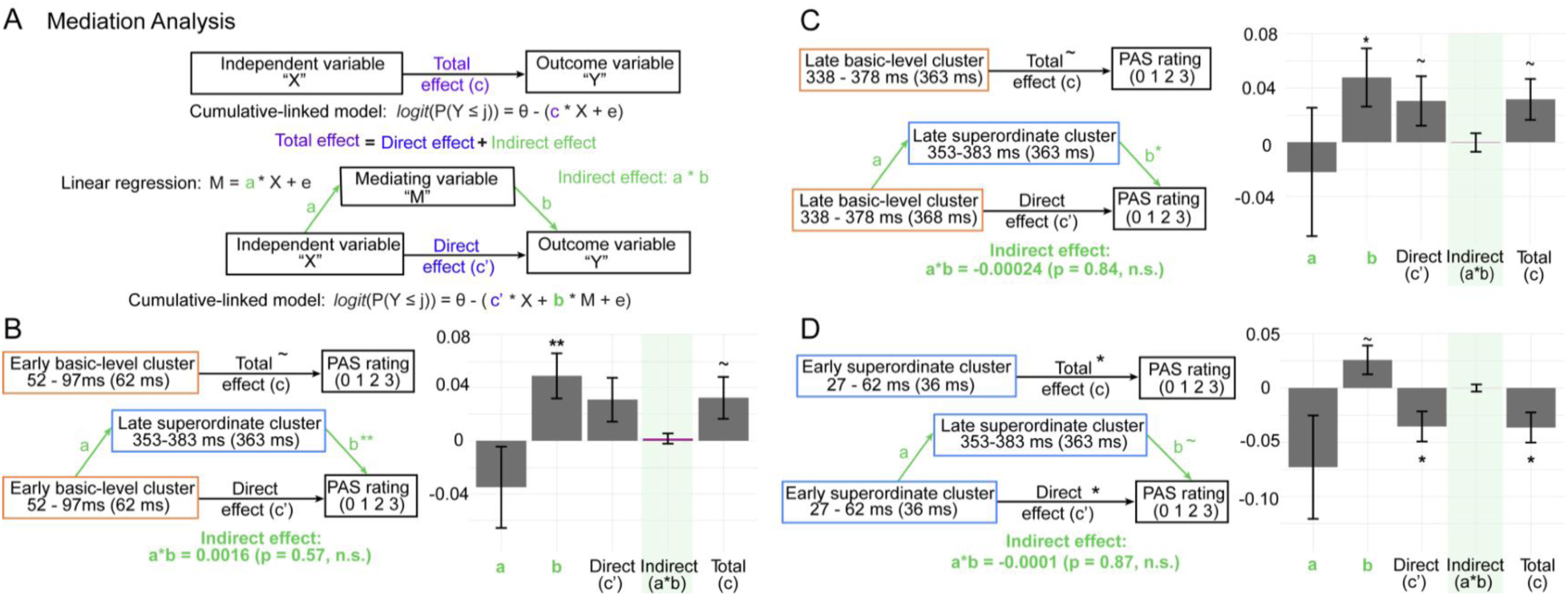
Mediation analysis. **(A)** Analysis schematic. The total effect of the independent variable (signed distance for a particular category classifier) on PAS ratings (path **c**) is decomposed into direct and indirect pathways via a mediating variable. Path **a** represents the relationship between the independent variable “X” and the mediator “M” with a linear regression model, and path **b** represents the relationship between the mediator “M” and the outcome “Y” via a cumulative-linked model. The direct effect is denoted by **c’,** and the indirect effect is quantified as the product **a*b**. (**B**) *Left*: Mediation analysis between the early basic-level cluster and the late superordinate cluster. For each independent variable, the time period for the significant cluster is followed by the peak timepoint in parentheses; classifier distance at the peak time point was input into the mediation model. *Right*: Group-level mean and s.e.m. of the coefficients obtained from the mediation model. Error bars represent the s.e.m. across subjects (n = 30). Statistics were evaluated by group-level Wilcoxon signed-rank test against 0. **(C-D)** Same as B, except that the early basic-level category representation is replaced by the late basic-level category representation (**C**) or the early superordinate category representation (**D**). ∼ p < 0.08 (marginal effects), * p < 0.05, ** p < 0.01.

First, we tested whether the late abstract superordinate category representation mediated the effect of the early or the late basic category representations on PAS ratings (given that all three neural metrics positively correlate with the PAS rating). To this end, for each participant, we identified the peak distance-to-bound value within each of the three significant clusters positively associated with PAS rating: early basic (peak latency: 62 ms), late basic (368 ms), and late superordinate (363 ms). We then evaluated two potential mediation models. In the first mediation model, we tested the effect of early basic-level category distance (independent variable “X”) on PAS ratings (outcome variable “Y”), mediated by the late superordinate category distance (mediation variable “M”) (**Fig. 6B**). In the second mediation model, we replaced the independent variable with the basic-level category distance from the second, late-stage positive cluster (**Fig. 6C**). For both models, the indirect effect was not significant (p = 0.57 for model 1, **Fig. 6B**, and p = 0.84 for model 2, **Fig. 6C**). The absence of mediation in both models suggests that the basic-level and superordinate category representations independently contribute to the subjective perceptual experience.

In model 3, we tested whether the late abstract superordinate category representation mediated the effect of the early superordinate category representation’s effect on PAS ratings. Although these two clusters have opposite relationships to PAS ratings (positive vs. negative), the later superordinate representation could, in principle, mediate the early superordinate representation’s behavioral influence. However, the indirect effect in this model was also not significant (p = 0.87, **Fig. 6D**), suggesting that conscious processing of objects does not proceed from an unconscious superordinate category representation to a conscious superordinate one; rather, these two category representations contribute independently, and could selectively emerge depending on the awareness level of that trial. In other words, a late abstract superordinate representation emerges on trials with high levels of awareness, and an early independent superordinate representation tends to occur on trials with poor awareness.

Together, these results demonstrate that basic-level and superordinate category representations independently contribute to perceptual awareness. Basic category representation correlates positively with perceptual awareness during both early and late time periods, and the late abstract superordinate category representation also correlates positively with awareness. By contrast, an independent early superordinate category representation is associated with poor perceptual awareness. Below, we discuss the interpretation and implication of these findings.

## DISCUSSION

In sum, by directly manipulating and measuring subjective awareness during object perception, we demonstrate that awareness has opposite relationships with early basic-level and early superordinate category representations: whereas the former is associated with more vivid perceptual experiences, the latter is associated with poor or absent awareness. Thus, although an early superordinate category representation exists, it does not contribute to perceptual awareness; however, we found that both superordinate and basic category representations positively contribute to objective categorization accuracy at the early stage. By contrast, a late superordinate category representation following an abstract format positively correlates with perceptual awareness. Therefore, the sequence of neural processing directly underlying conscious object perception proceeds from the basic level to the superordinate level.

Our observation of an early unconscious superordinate category representation is supported by several convergent findings: *First*, superordinate, but not basic-level, category information is significantly decoded at 160–280 ms on unconscious (PAS = 0) trials (**Fig. 3G**). *Second*, the strength of early superordinate category representation *negatively* correlates with PAS ratings from trial to trial (**Fig. 5E**). *Third*, behaviorally, superordinate categorization performance is significantly above the chance-level in unaware (PAS = 0) trials in the LSF condition (**Figs. 2F**, **S2E**). Crucially, these findings reconcile previously conflicting observations supporting, alternatively, a coarse-to-fine sequence versus a fine-to-coarse sequence of processing by demonstrating that an early superordinate (coarse) representation indeed exists in neural activity, but it does not directly contribute to awareness. In fact, our observation of a *negative* association between this neural representation and awareness suggests that it might compete with the neural processing that directly underlies awareness — a possibility we elaborate on below.

By contrast, we observed that an early basic-level category representation positively correlates with perceptual awareness (**Fig. 5E**). This finding significantly extends previous reports of an early emergence of basic-level category representation in the neural activity (Carlson et al., 2013; Greene and Rohan, 2025) by establishing that this activity indeed contributes to perceptual awareness. This early onset time is consistent with a feedforward sweep in the ventral visual stream, which reaches IT cortex within 100 ms (Rousselet et al., 2004).

In addition, we demonstrate that a late-stage, abstract superordinate category representation positively correlates with perceptual awareness. The abstract nature of this superordinate category representation is confirmed by the neural geometry analysis, which shows that the superordinate category is represented in a format disentangled from the idiosyncratic features unique to each basic category, such that a decoder trained on one pair of basic categories can generalize to a distinct pair of basic categories that it has never seen during training. In support of this interpretation, a previous study reported that animacy decoding using texform stimuli reached peak decoding accuracy within 200 ms (Wang et al., 2022)—a distinct temporal profile as compared to our CCGP decoding analysis, which slowly increased over time (Fig. 4B-D). Lastly, although our stimulus set is relatively small, consisting of 12 exemplars, previous studies on abstract neural representations have used stimulus sets of comparable sizes (Courellis et al., 2024; Liang et al., 2026; Wójcik et al., 2026); this design choice allows a larger number of trials to be collected for each exemplar. Together, these results suggest that object category processing underlying perceptual awareness proceeds from an early basic-level category representation to a late, abstract superordinate category representation.

Our neural findings also help to reconcile a previous debate in the behavioral literature. Although the basic category advantage is well-established in behavioral studies (Rosch et al., 1976; Jolicoeur et al., 1984), a few studies have shown that reaction times for superordinate category discrimination can be faster than basic category discrimination (Mace et al., 2009; Loschky and Larson, 2010; Prass et al., 2013). Typically, studies reporting a basic-level advantage have used a free-naming style task, whereas studies reporting a superordinate advantage have used a force-choice discrimination task. Classic neuropsychology work (Milner and Goodale, 2008) suggests that naming/identification tasks rely more on the conscious route, whereas discrimination performances can be driven by unconscious processing. Thus, these divergent behavioral findings can be explained by our present observation that the basic-level representation directly contributes to perceptual awareness, whereas an early superordinate representation can drive successful discrimination behavior unconsciously.

Our observation that superordinate categorization behavior on unconscious trials is significantly above-chance in the LSF condition is notable. Given that the unconscious superordinate neural representation occurs quite early (**Fig. 5E**), these results are broadly consistent with the idea that LSF information tends to be processed faster in the brain due to its preferential processing by the magnocellular visual pathway (Merigan and Maunsell, 1993; Kveraga et al., 2007; Schuurmans et al., 2023). This early superordinate representation might rely on relatively coarse visual cues, such as the rounded curvatures of animate objects and the sharp edges and lines of manmade objects (Long et al., 2018; Wang et al., 2022). Speculatively, early superordinate category representations that emerge despite poor or absent awareness might be evolutionarily advantageous for survival, by triggering appropriate action circuits prior to full awareness.

We note that “unconscious perception”—above-chance objective performance in the absence of conscious awareness—is sometimes contested in the literature, because of a potential confound of decision biases (Peters and Lau, 2015; Phillips, 2021; Michel, 2023). However, alternative approaches such as using chance-level objective performance as an indication of the absence of awareness, run into caveats with potentially significant underestimation of unconscious processing since unconscious processing can influence behavioral performance, as demonstrated by subliminal priming (e.g., (Dehaene et al., 1998; van Gaal et al., 2008). In our study, the validity of our subjective report claims was strengthened by using a 4-point perceptual awareness scale (PAS) rating, which captures subjective experiences more precisely than binary responses (seen/not seen) or confidence ratings (Overgaard et al., 2006; Sandberg et al., 2010; Overgaard and Sandberg, 2021). In particular, by using strictly PAS = 0 trials (“not even the faintest sensation) for the definition of “unconscious” trials, we exclude the unclear/borderline cases of weak conscious perception associated with PAS = 1 (“a glimpse”). Although the use of PAS ratings does not remove the decision bias entirely, the flexibility to report graded experiences significantly removes potential ambiguity surrounding subjective reports.

Motivated by the observation that the early basic-level category representation and the early superordinate category representation have opposing relationships to perceptual awareness, we propose a mechanistic, ‘dual-stream’ model to account for these findings. Under this model, the early superordinate category representation is mediated by a fast, largely unconscious feedforward sweep through the dorsal visual stream reaching the frontal cortex (**Fig. 7**, left), whereas the early basic category representation is mediated by feedforward processing in the ventral visual stream (**Fig. 7**, right). In addition, we postulate that the latter pathway, through recurrent processing in the ventral stream and between the ventral stream and downstream prefrontal regions, eventually leads to the abstract superordinate category representation at the late stage. This proposal is supported by our channel-level searchlight decoding analysis showing that both the early superordinate category representation and the late abstract superordinate category representation have frontal contributions (**Fig. 3D-F** and **Fig. S8**). It also resonates with an early theory proposing that fast, low-spatial-frequency information could rapidly reach the frontal cortex via the dorsal pathway, the result of which could guide ventral stream processing (Bar et al., 2006; Bar, 2007). Of note, a recent non-human primate neurophysiology study provides direct evidence for this theory, demonstrating that the LSF information of images is captured in *early-stage* ventrolateral prefrontal cortex (vlPFC) activity, 50–90ms after image onset, and supports coarse category representations (Bellet, 2026). In accordance, earlier works show that vlPFC encodes visual content especially when there is stimulus ambiguity (Wang et al., 2013; Kar and DiCarlo, 2021; Kapoor et al., 2022; He, 2023; Panagiotaropoulos, 2024). Importantly, our present results suggest that this initial fast processing is largely unconscious.

**Figure 7.**
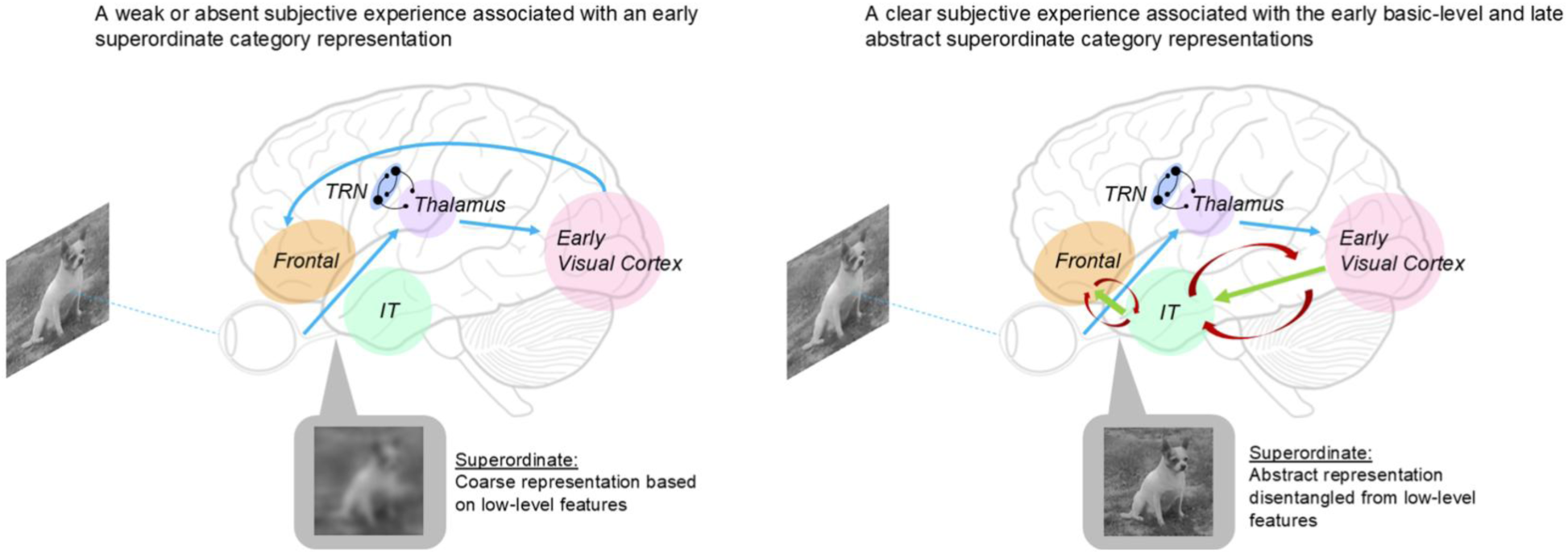
A hypothesized dual-stream model to account for our neural findings. During low awareness trials, stimulus processing is biased to the dorsal stream by the lateral inhibition circuit of the thalamic reticular nucleus (TRN), resulting in a coarse supraordinate representation. By contrast, a cascading feedforward sweep (green arrow) and recurrence (red arrows) through the ventral visual stream to the inferotemporal (IT) cortex and frontal cortex results in a clear subjective experience of the object, and supports both an early basic-level category representation and a late abstract superordinate category representation via disentanglement of low-level object features.

What could cause the apparent competitive relationship between these two pathways? We hypothesize that the thalamus might play a key role here. First, the pulvinar, a higher-order thalamic nucleus, plays an integrative role in visual processing and modulates both feedforward and feedback visual processing pathways (Saalmann and Kastner, 2011; Cortes et al., 2024). A retina-pulvinar-dorsal pathway has been proposed as a visual pathway underlying blindsight (Warner et al., 2015). Second, the lateral inhibition mechanism within the thalamic reticular nucleus (TRN), in concert with its reciprocal connection to the pulvinar, might support the apparent competitive mechanism between early superordinate and early basic-level category representations observed here. The TRN is a thin layer of GABAergic neurons and receives inputs from both the thalamus and the cortex (Pinault, 2004; Halassa and Acsády, 2016). We speculate that during feedforward processing of retinal inputs, the lateral inhibition within the TRN might control its gating of the pulvinar nucleus and biases information processing in favor of one of the two visual pathways. In support of this idea, TRN neurons selectively inhibit different thalamic nuclei and show topographic and patchy patterns with preferential targeting of specific pulvinar subdivisions (FitzGibbon et al., 1995); additionally, different subdivisions of the pulvinar preferentially connect to the dorsal stream or the ventral stream (Kaas and Lyon, 2007; Saalmann and Kastner, 2011; Bilderbeek et al., 2026). More broadly, the thalamocortical circuit is well-suited to modulating perceptual awareness and attention (Saalmann et al., 2012; Whyte et al., 2024). As such, the anatomical infrastructure for supporting our proposed dual-stream model exists. However, this mechanistic model remains to be tested by future studies employing techniques with finer resolutions and subcortical access (e.g., intracranial EEG) and/or causal manipulation.

Importantly, our findings suggest that an influential theory proposing a coarse-to-fine sequence of processing underlying conscious vision (Hochstein, 2002) needs to be updated. This theory suggests that “conscious perception begins at the hierarchy’s top” and that the initial conscious percept is “vision at a glance”, followed by “vision with scrutiny”. Our results do not dispute the existence of an early, coarse, “gist-like” representation, which affords superordinate category representation, but we demonstrate that this early coarse representation is unconscious — in fact, the stronger this representation, the weaker the reported perceptual awareness.

We note that most of the behavioral studies that initially motivated the coarse-to-fine theory of conscious vision were based on visual search or perceptual learning paradigms where the effects had key contributions by attention or plasticity. Our findings do not challenge these earlier empirical observations; rather, they demonstrate that a straightforward application of this coarse-to-fine idea to conscious vision as previously proposed (Hochstein, 2002) does not stand. Our findings suggest a revised theoretical framework: conscious object perception starts at the mid-level (basic category) and proceeds to both higher (abstract superordinate category) and lower (detailed visual features) levels through feedforward and feedback connections, respectively. We look forward to future empirical tests of this idea. If proven, this framework has important implications for theories of conscious vision, as it is only partially compatible with prominent theories such as the Global Neuronal Workspace theory (Mashour et al., 2020) or Recurrent Processing Theory (Lamme, 2006). This is not surprising given that current theories leave many questions unaddressed (He, 2023, 2025).

Interestingly, recent rodent and primate studies have reported coarse-to-fine receptive field dynamics in V1 neurons, where neurons respond first to low-spatial-frequency information and then to high-spatial-frequency information (Skyberg et al., 2022; Parker et al., 2023). We believe that there is no tension between these findings and our findings, and indeed, the early coarse information represented in V1 might be related to the early unconscious superordinate representation we observed. An important question for future investigation is whether the sequence of category-level processing underlying conscious object perception revealed herein remains the same during free eye movements, when saccades introduce temporal modulation to the spatial information in the visual inputs received by the retina (Boi et al., 2017).

In conclusion, we reveal that conscious object perception proceeds from a basic category representation to an abstract superordinate category representation; in parallel, an early superordinate category representation is largely unconscious and appears to compete with the conscious processing. These results offer new insights into the sequence of category-level neural processing underlying conscious vision.

## Data and Code Availability

The code and source data to reproduce all figures will be provided with the published paper.

## Acknowledgements

This work was supported by a grant from U.S. National Institutes of Health (R01EY032085, to BJH), as well as NYU Grossman School of Medicine. TJB acknowledges support by European Union’s Horizon 2020 research and innovation programme under the Marie Skłodowska-Curie grant agreement No. 795998. We would like to thank Shira Baror, Richard Hardstone and Michael Zhu for technical assistance and helpful discussions.

