## Supplementary Information for "Probing the neural geometry of conscious object perception in time"

Hachisuka, Baumgarten, et. al.

Including:

**Supplementary Results**

**Supplementary Methods**

**9 Supplementary Figures**

### SUPPLEMENTARY RESULTS

#### Experiment 1: Validating the object-substitution orientation-shifted mask (OSOM), part 1.

We developed a novel visual masking approach, called Object Substitution Orientation-Shifted Mask (OSOM). Similar to the classic object substitution mask (OSM) (Koivisto et al., 2014; Koivisto and Kahila, 2017), the target and mask appear on the screen at the same time ("common onset"), followed by the mask staying on the screen for an additional amount of time ("delayed offset"). The mask consists of dots that surround the corners of the target image. OSM is thought to selectively impair recurrent processing more precisely, compared to classical visual masking approaches. For example, unlike backwards masking, the target and mask do not spatially overlap. This keeps the feedforward processing intact, while more selectively disrupting the recurrent processing (Carlson et al., 2013). Additionally, unlike metacontrast masking (Del Cul et al., 2007), the mask does not follow the contour of the target image. This mitigates the confound that lateral inhibition between the target and mask during the feedforward sweep, driven by shared contours, could contribute to visibility impairments of the target (Breitmeyer, 2015). However, traditional OSM produces relatively weak perceptual impairments. Thus, to strengthen the mask's effect, we developed a modified OSM paradigm, called OSOM (object substitution orientation-shifted mask). In this mask, we rotated the *trailing* mask by 90° relative to its initial orientation at the onset.

For either OSM or OSOM, spatial attention needs to be dispersed for the mask to be effective; to facilitate this condition, the target-mask pair was presented in 1 of 4 peripheral locations with equal probability (top-right, top-left, bottom-right, or bottom-left) (Koivisto et al., 2014; Koivisto and Kahila, 2017). Phase-scrambled filler images were displayed in the other three locations.

The two experiments described below were conducted to verify the behavioral effectiveness of OSOM masking and were conducted in separate participant groups as compared to the main experiment. Each experiment began with an adaptive staircase procedure to titrate each exemplar's image contrast in the normal image condition to a ~63% basic-level categorization performance. Subsequently, the same image contrast was used in all image conditions.

The trial structure was as follows, mirroring the paradigm structure in the main text: following each image presentation, participants answered two alternative forced-choice (2AFC) questions regarding the perceived object category: first for the superordinate category ("Animal" or Vehicle") and then for the basic category ("Dog" or "Cat" if they answered "Animal", "Car" or "Truck" if they answered "Vehicle") (**Supplementary Fig. 1A, 2A**). If the participant responded with an incorrect superordinate category, invalid basic-category options were displayed to be consistent with their initial answer. In all subsequent analyses, basic-level categorization performance is computed only from correct superordinate categorization trials, thereby excluding trials with invalid basic-category choices.

In Experiment 1, there were three possible image conditions: Normal, OSM, and OSOM. We found that OSOM is a stronger mask than OSM. A two-way repeated measures ANOVA [factors: category level (Superordinate and Basic) × condition (Normal, OSM, and OSOM)] (**Supplementary Fig. 1B-C)** revealed a non-significant interaction effect ($F_{1,15}$ = 0.03, p = 0.97, $\eta_p^2$ = 0.002), suggesting that image conditions similarly affect both categorization performances. The main effects of image condition ($F_{1,15}$ = 71.4, p = 3.96e-12, $\eta_p^2$ = 0.83) and category level ($F_{1,15}$ = 105.6, p =3.47e-8, $\eta_p^2$ = 0.88) were both significant, suggesting that categorization performance was perturbed by the image manipulations, and that the superordinate categorization performance is more robust than that of the basic level. The Shapiro-Wilk test did not indicate a significant deviation from normality (W = 0.985, p = 0.345). Post hoc tests using the Bonferroni correction revealed significant decreases in categorization performance across image conditions. Compared with the Normal condition, OSOM most strongly decreased both superordinate (p

= 0.002) and basic-level ($p$ = 3.45e-8) categorization performance. Compared to OSM, OSOM showed a marginal but non-significant decrease in superordinate categorization performance ($p$ = 0.056) and a significant decrease in basic categorization performance ($p$ = 0.028). OSM images also showed decreased performance compared to the normal condition (Superordinate: $p$ = 0.002; Basic: $p$ = 0.006).

Using a one-way repeated-measures ANOVA, we also evaluated the effect of image condition (Normal, OSM, and OSOM) on PAS ratings (**Supplementary Fig. 1D**), which revealed a significant main effect of condition ($F_{1,15}$ = 39.9, $p$ = 3.52e-9, $\eta_p^2$ = 0.73). The Shapiro-Wilk test did not indicate a significant deviation from normality ($W$ = 0.960, $p$ = 0.09). Post-hoc pairwise comparisons using the Bonferroni correction showed that the PAS ratings for OSM were significantly reduced compared to Normal ($p$ = 2.36e-5). Moreover, PAS ratings for OSOM were significantly reduced compared to Normal ($p$ = 3.42e-9) and OSM images ($p$ = 0.003). Together, these results suggest that OSOM is a stronger mask than the classic OSM mask.

Why is OSOM more effective at impairing visual processing than OSM? Below, we investigate this question in an additional behavioral experiment.

**Experiment 2: Validating OSOM, part 2.**

One possibility is that the enhanced effect of the OSOM could simply reflect an increased salience from the orientation change, rather than a stronger disruption of recurrent processing. This would imply that object visibility failure is caused by a shift in exogenous attention rather than by a disruption in perceptual processing. To assess this possibility, in Experiment 2, we moved the trailing mask to a new location while maintaining its orientation shift (**Supplementary Fig. 2A**). This manipulation preserves the salience of the mask's orientation change but prevents the target and mask from forming a unified object representation (Lleras and Moore, 2003). Below, we compared OSM, "OSOM-stable" (OSOM from Experiment 1), and the control OSOM mask called "OSOM-moved."

A two-way repeated measures ANOVA [factors: category level (Superordinate and Basic) $\times$ condition (OSM, OSOM-stable, and OSOM-moved)] to assess image condition's effect on categorization performance (**Supplementary Fig. 2B-C**). This revealed a significant interaction effect ($F_{1,8}$ = 5.5, $p$ = 0.025, $\eta_p^2$ = 0.41), as well as significant main effects of the image condition ($F_{1,8}$ = 24.6, $p$ = 1.31e-5, $\eta_p^2$ = 0.76) and category level ($F_{1,8}$ = 91.2, $p$ = 1.20e-5, $\eta_p^2$ = 0.92). The Shapiro-Wilk test did not indicate a significant deviation from normality ($W$ = 0.984, $p$ = 0.703). Post-hoc tests using Bonferroni correction revealed significant decreases in superordinate categorization performances for OSOM-stable compared to OSM ($p$ = 1.37e-4) and for OSOM-stable compared to OSOM-moved ($p$ = 5.35e-7). A similar pattern was observed for basic-level categorization performances, with significant differences between OSOM-stable and OSOM-moved ($p$ = 0.048), though not between OSOM-stable and OSM ($p$ = 0.346). There were no significant differences between OSM and OSOM-moved categorization performances (Superordinate: $p$ = 0.86, Basic: $p$ = 1).

Using a one-way repeated-measures ANOVA, we also evaluated the effect of image condition (OSM, OSOM-stable, and OSOM-moved) on PAS ratings (**Supplementary Fig. 2D**). This revealed a significant main effect of image condition ($F_{1,8}$ = 11.1, $p$ = 9.34e-4, $\eta_p^2$ = 0.58). The Shapiro-Wilk test did indicate a significant deviation from normality ($W$ = 0.8774, $p$ = 0.004). However, a non-parametric test using the Friedman rank sum test confirmed a significant main effect of the image condition on PAS ratings ($\chi^2(2)$ = 9.56, $p$ = 0.008). Post-hoc pairwise comparisons using the Bonferroni correction showed that the PAS ratings for OSOM-stable were significantly reduced compared to OSM ($p$ = 0.01) and OSOM-moved ($p$ = 0.001). There was no significant difference between OSM and OSOM-moved ($p$ = 0.30).

Together, these results show that our OSOM manipulation ('OSOM-stable' in Experiment 2) causes the largest impairment in both categorization performance and awareness ratings, despite being a less attention-grabbing mask than the OSOM-moved condition, where the mask is both rotated and moved to a different location. This suggests that the reduction in target visibility in the OSOM condition, as compared to the classic OSM mask is achieved by perturbing recurrent processing, rather than by causing an exogenous attention shift. Specifically, according to the object updating theory of OSM (Goodhew et al., 2013; Goodhew et al., 2015; Goodhew, 2017), the target and the mask are bound together in time, which results in the target being dropped from perceptual representation due to the trailing mask. The rotated mask might strengthen this effect due to triggering stronger recurrent processing related to the mask processing (Lleras and Moore, 2003).

**SUPPLEMENTARY METHODS — related to Supplementary Results and Supplementary Figs. 1-2**

**Participants**

Participants were recruited from the greater New York City area. All participants provided written informed consent, and the experiment was approved by the Institutional Review Board of New York University Grossman School of Medicine (protocol #s15-01323).

19 participants were recruited for Experiment 1 (11 male, 8 female, mean age: 26 years) and 12 participants were recruited for Experiment 2 (7 male, 5 female, mean age: 23 years). All participants had normal or corrected-to-normal vision and were right-handed except for one participant in Experiment 1.

3 participants from Experiment 1 and 2 participants from Experiment 2 were excluded after exhibiting poor staircase behavior. Poor staircase behavior was defined as failure to reach a constant threshold on 3 or more exemplar images. Additionally, 1 participant from Experiment 1 and 1 participant from Experiment 2 were excluded because they had chance-level basic categorization performance for normal image conditions during the main task, and 1 participant from Experiment 2 was excluded for skipping more than half of the trials in the main task. In total, data from 16 participants in Experiment 1 (9 male, 7 female, mean age: 24 years) and 9 participants in Experiment 2 (6 male, 3 female, mean age: 23 years) were used for analysis.

**Behavioral Task Design**

*Stimuli*

The image set is the same as in the main task. Stimuli size was 2.4 dva, and all targets appeared pseudo-randomly in the periphery at 5.01 dva from fixation (top-right/top-left/bottom-right/bottom-left). The 4 dots around the target were each 1.2 dva in diameter, and each dot was placed at 0.27 dva from each corner of the target image. As in the main task, the remaining three locations had phase-scrambled images created from original images that fell in the other three basic categories as compared to the target image.

*Experiment 1 image manipulations*

In Experiment 1, the following image manipulations were used: Normal (unaltered images), Object Substitution Mask (OSM), and the novel, Object Substitution Orientation-Shifted Mask (OSOM). The only difference between OSM and OSOM was the orientation of the trailing mask. In OSM, the trailing mask kept the same orientation of the mask at the target image onset, at the corners of the target image. In OSOM, the trailing dots are rotated 90° relative to the original orientation.

*Experiment 2 image manipulations*

In Experiment 2, the following image stimulus conditions were used: Object Substitution Mask (OSM), the novel OSOM condition with the rotated trailing mask (“OSOM-stable”, identical to OSOM in Experiment 1), and a modified OSOM with a position-shifted trailing mask (“OSOM-moved”). In OSOM-moved, the trailing mask was still rotated 90°, but additionally, its position was pseudo-randomly determined, ensuring it did not overlap with the initial target and mask location.

*Staircase*

The staircase procedure is identical to that of the main task; see main text for details.

*Task*

Both experiments consisted of 576 trials in total, evenly distributed across the 3 image conditions (192 trials/condition). In each session, trials were subdivided into 3 blocks, with 192 trials per block. Unlike the main task, this session included 435 target-image trials and 141 catch trials, each evenly distributed across the session. In the catch trials, all 4 locations had phase-scrambled images, with the mask presented at one of the conditions (except for the normal condition in Experiment 1). However, for consistency with the analysis in the main text, only the behavioral results from the target image presentations are analyzed here.

*Behavioral analysis*

Behavioral analysis follows the same methods as in the main experiment. Briefly, for all behavioral analyses, the basic-level categorization performance is computed only on correct superordinate categorization trials. Repeated measures ANOVA was conducted using Jeffrey’s Amazing Statistics Program (JASP) (Team, 2024). Normality assumption of data distribution was checked using the Q-Q plots of the model residuals using the “afex” package in R, and the Shapiro-Wilk test. When a deviation from normality was detected, a non-parametric test using the Friedman rank sum test was performed.

For all analyses, post hoc comparisons were conducted to examine significant main effects. P-values for post-hoc analyses were adjusted using the Bonferroni correction for multiple comparisons. For the post-hoc analyses for the main effect of image condition on PAS ratings in Experiment 2, a Wilcoxon signed-rank test with Bonferroni correction was performed.

**References** are included in the bibliography in the main text.

### SUPPLEMENTARY FIGURES

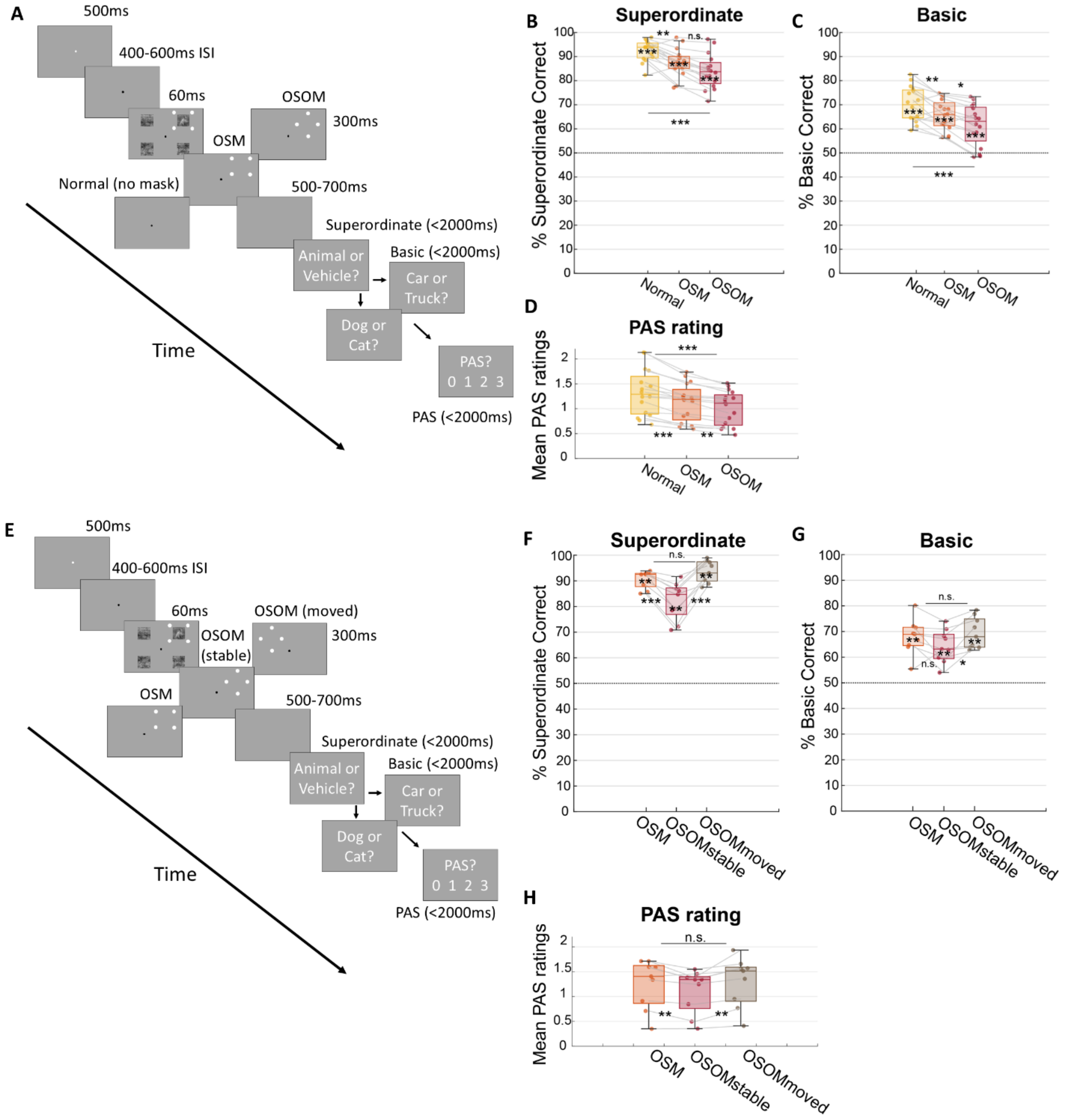


**Supplementary Figure 1: [Top] Object Substitution Orientation-shifted Mask (OSOM) is more effective than the Object Substitution Mask (OSM). (A)** Trial structure. Images were presented for 60 ms in one of three image conditions (Normal, OSM, and OSOM), followed by two categorization questions and a subjective vividness report using the perceptual awareness scale (PAS) ratings from 0 (“no experience”) to 3 (“clear experience”). The target image location was pseudo-randomly chosen between top-right, bottom-right, bottom-left, and top-left locations with equal probability, and filler images (phase-scrambled images) were placed in the remaining slots. **(B)** Superordinate categorization performance for each image condition. The box plots show the median (middle line) and the lower/upper quartiles (top and bottom of the box) across participants (n = 16). Light gray lines connect performances for the same participant across conditions. **(C)** Same as (B) but for basic-level categorization responses. Only correct superordinate trials (which prompt valid basic categorization questions) are included. In B-C, chance-level accuracy is at 50%. **(D)** Mean PAS ratings across three image conditions. Format is the same as B-C. Asterisks between box plots indicate significance based on Bonferroni-corrected post-hoc statistics. **[Bottom] Object Substitution Orientation-shifted Mask (OSOM) relies on temporal integration instead of attentional distraction. (E)** Trial structure. Images were presented for 60 ms in one of three image conditions (OSM, OSOM-stable, and OSOM-moved), followed by two categorization questions and a subjective vividness report. The target image location was randomly chosen between top-right, bottom-right, bottom-left, and top-left with equal probability, and filler images (phase-scrambled images) were placed in the remaining slots. **(F)** Superordinate categorization performances across the three image conditions. The box plots show the median (middle line) as well as the lower and upper quartiles (top and bottom of the box). Light gray lines connect performances for the same participant across conditions ($n = 9$). **(G)** Same as (B) but for basic-level category responses. Only correct superordinate trials (that prompt valid basic category questions) are included. **(H)** Mean PAS ratings across three image conditions. Asterisks between box plots indicate significance based on Bonferroni-corrected. *: $p < 0.05$, **: $p < 0.01$, ***: $p < 0.001$.

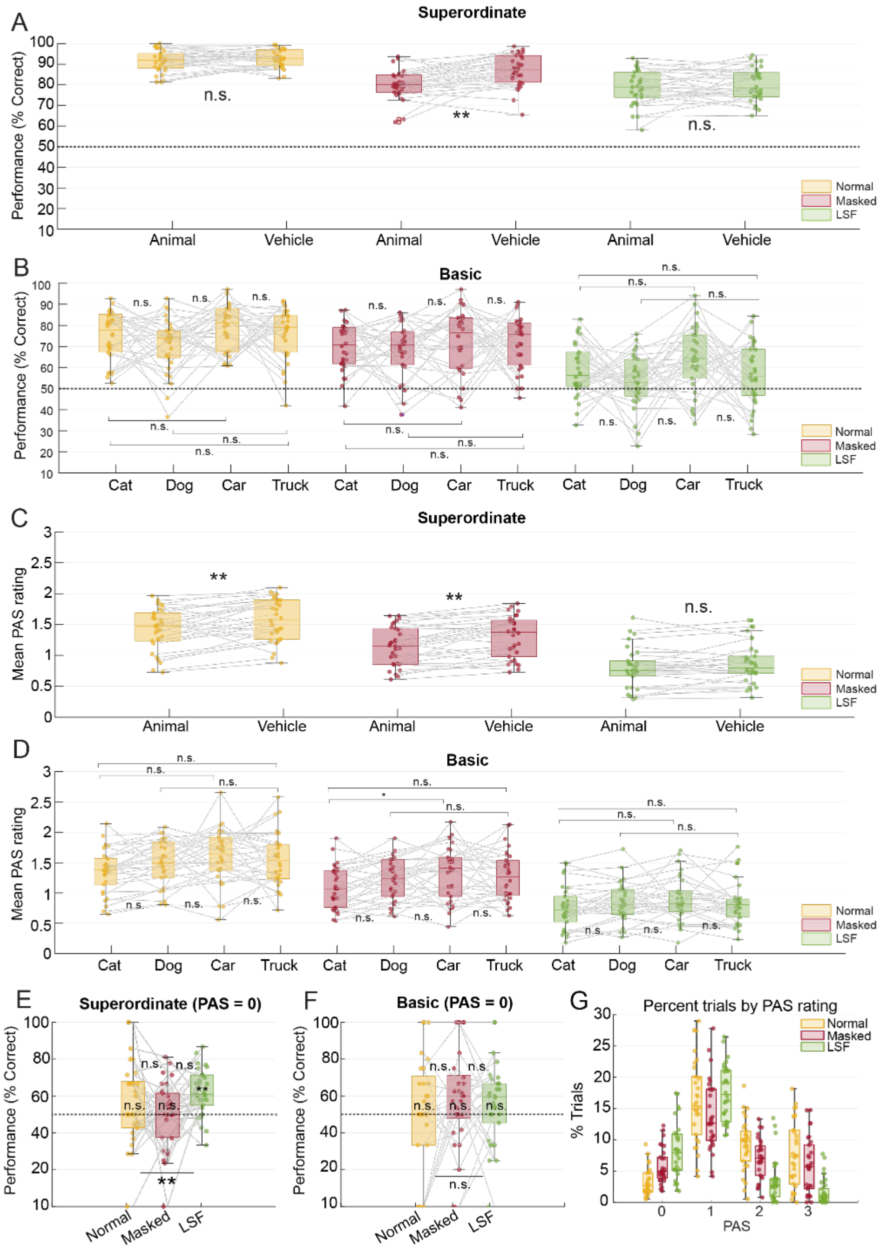


**Supplementary Figure 2. Detailed analyses on categorization performance from the main task (n = 30). (A)** Categorization performance broken down by superordinate categories (animal vs. vehicle). The box plots show the median (middle line) as well as the lower and upper quartiles (top and bottom of the box). Each dot represents an individual subject, and the light gray lines connect performance of the same individual across conditions. **(B)** Categorization performance broken down by basic-level categories (cat vs. dog vs. car vs. truck). **(C)** PAS ratings broken down by superordinate categories (animal vs. vehicle). **(D)** PAS ratings broken down by basic-level categories. **(E)** Superordinate categorization performances across the three image conditions in PAS = 0 (undetected) trials only. **(F)** Same as (E) but for basic-level categorization responses. Only correct superordinate trials (that prompt valid basic category questions) are included. Asterisks between box plots indicate significance based on Bonferroni-corrected post-hoc statistics from a 2-way repeated measures ANOVA. Asterisks printed on top of the box plots are statistics from Wilcoxon signed-rank tests against 50% chance-level performance (FDR-corrected). *: $p < 0.05$, **: $p < 0.01$, ***: $p < 0.001$. **(G)** Proportion of total trials for each PAS rating response, plotted separately for each image condition.

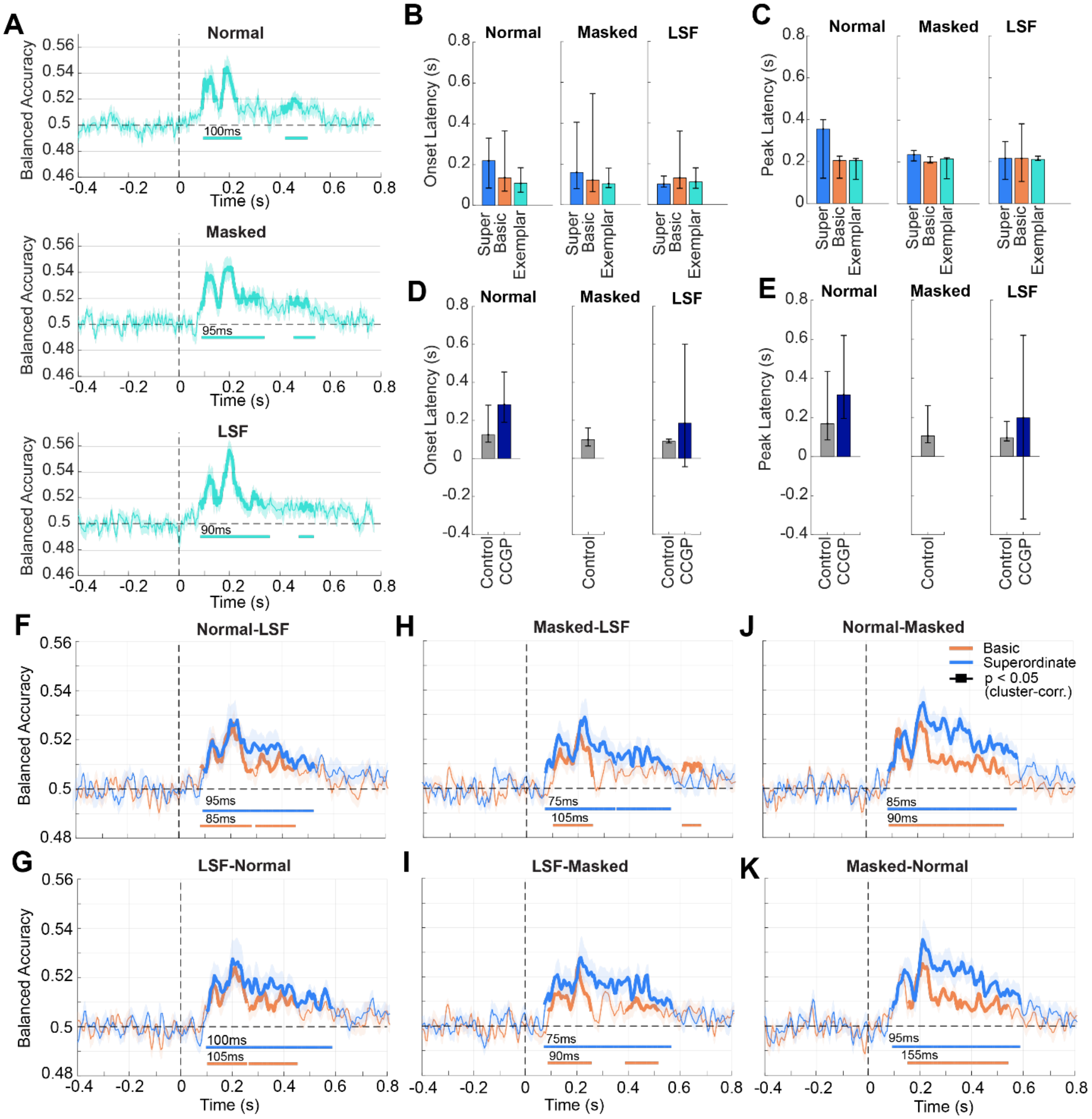


**Supplementary Figure 3: Additional analyses (A)** Average decoding accuracy traces for discriminating exemplar images. Classifiers were trained to discriminate between exemplar images within each basic level category: dog 1 vs. dog 2 vs. dog 3; cat 1 vs. cat 2 vs. cat 3; car 1 vs. car 2 vs. car 3; truck 1 vs. truck 2 vs. truck 3. The traces indicate decoding analysis results from Normal (top), Masked (middle), and LSF (bottom) image conditions. Shaded areas indicate the s.e.m. across subjects (n = 30). Bolded traces and horizontal lines indicate significant timepoints ($p < 0.05$, cluster-based permutation test). Numbers indicate the onset times of each statistically significant time segment. **(B-C)** Group-level onset times (**B**) and peak latency times (**C**) for Fig. 3A-C (n = 30). **(D-E)** Group-level onset times (**D**) and peak latency times (**E**) for Fig. 4B-D. In all plots, bar height indicates group-level mean, and error bars indicate the 95% distribution of confidence intervals, derived from bootstrapping onset or peak times across subjects. **(F-K)** Average cross-condition decoding accuracy traces for superordinate (animal vs. vehicle, in blue) and basic category (dog vs. cat, or car vs. truck; within each superordinate category, in orange) classifications. The two decoding accuracy traces for basic category decoding were averaged within each subject. Within each subtitle, the first condition is the training condition, the second condition is the testing condition; e.g., panel F is training on the Normal condition and testing on the LSF condition. (G-I) are the same as Fig. 3I-J in the main text. Shaded areas indicate s.e.m across subjects (n = 30). Bolded traces and horizontal lines indicate significant timepoints ($p < 0.05$, cluster-based permutation test). Numbers indicate the onset times of each statistically significant time segment.

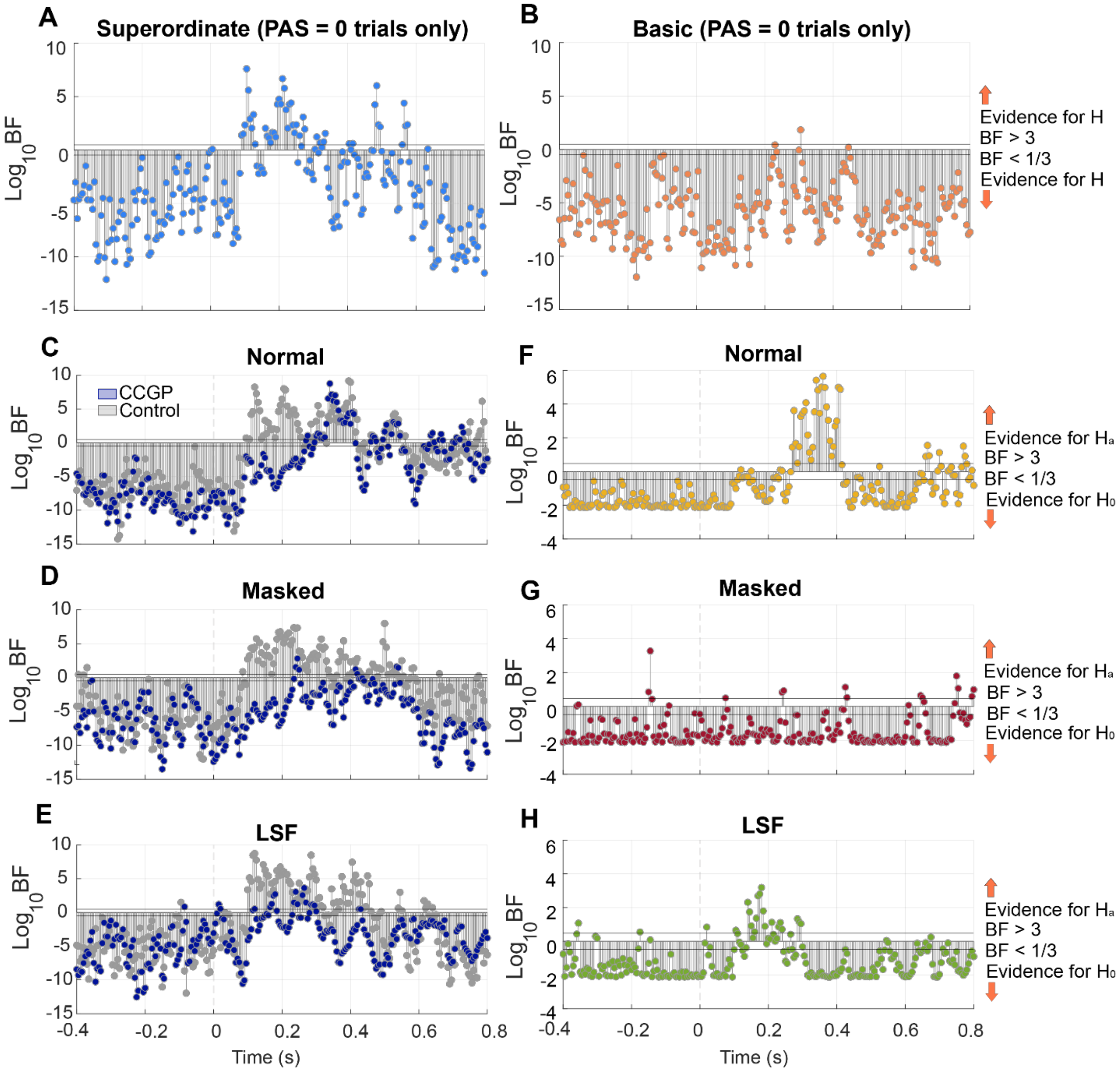


**Supplementary Figure 4: Bayesian statistics corresponding to multivariate pattern analyses. (A-B)** Bayes Factors for the multivariate pattern analysis for PAS = 0 trials, corresponding to Fig. 3G. Log-transformed Bayes Factors (BF) as a function of time for **(A)** superordinate (animal vs. vehicle) **(B)** basic (dog vs. cat; car vs. truck) category decoding analyses (n = 30). Timepoints with BF > 3 suggest evidence for the alternative hypothesis ($H_a$), whereas BF < 1/3 suggests evidence for the null hypothesis ($H_0$). BF = 3 and BF = 1/3 are indicated by the two horizontal lines flanking the y = 0 horizontal line. **(C-E)** Bayes Factors for the Cross-Category Generalization Performance (CCGP) analysis, corresponding to Fig. 4B-D. Plotted values are log-transformed Bayes Factors (Log

$BF_{10}$) plotted as a function of time (n = 30), corresponding to Bayesian statistics for the CCGP decoding accuracy traces in Fig. 4B-D. **(F-H)** Log-transformed Bayes Factors (Log $BF_{10}$) plotted as a function of time (n = 30), that correspond to the Parallelism Score analysis in Fig 4F. See Figure 4A for the CCGP analysis schematic, and Fig. 4E for the Parallelism Score analysis schematic. Timepoints with BF > 3 suggest evidence for the alternative hypothesis ($H_a$), whereas BF < 1/3 suggests evidence for the null hypothesis ($H_0$). BF = 3 and BF = 1/3 are indicated by the two horizontal lines flanking the y = 0 horizontal line.

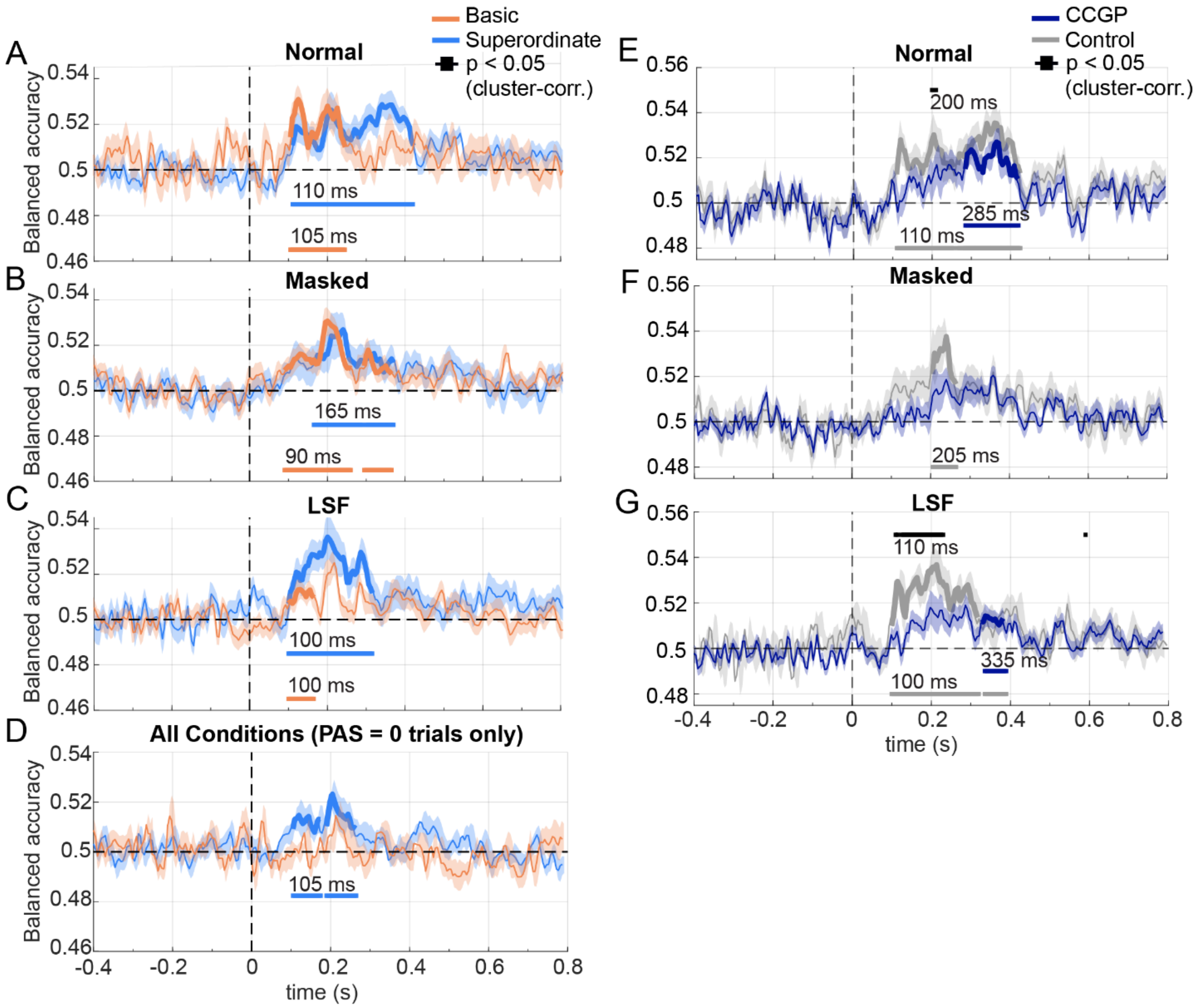


**Supplementary Figure 5: Superordinate, Basic, and CCGP decoding analysis using a cross-image-location decoding scheme.** Similar to Fig. 3 (A-C, G) and Fig. 4 (B-D) in the main text, but with classifiers trained on trials that did not share target image location as the left-out test trials. **(A-C)** Cross-image location decoding analysis results for superordinate (cat-and-dog vs. car-and-truck) and basic-level (average of cat vs dog and car vs. truck) categories in the Normal, Masked, and LSF conditions; corresponds to Fig. 3A-C in the main text. Classifiers are trained using trials from 3 target image locations (e.g., top/bottom/right) and tested on trials from the remaining target image location (e.g., left). Decoding accuracies were averaged across the four train-test splits. **(D)** Same as (A-C), but for trials with PAS ratings reported as “0” (no perceptual experience), corresponding to Fig. 3G. **(E-G)** Cross-image location CCGP and control analysis corresponding to Fig. 4B-D. In all panels, shaded areas indicate s.e.m. across subjects (*n* = 30). Bolded traces and horizontal lines indicate significant timepoints ($p < 0.05$, cluster-based permutation test). The black horizontal lines above the traces in E-G indicate significant differences between the CCGP and control decoding accuracies. Numbers indicate the onset times of each statistically significant time segment.

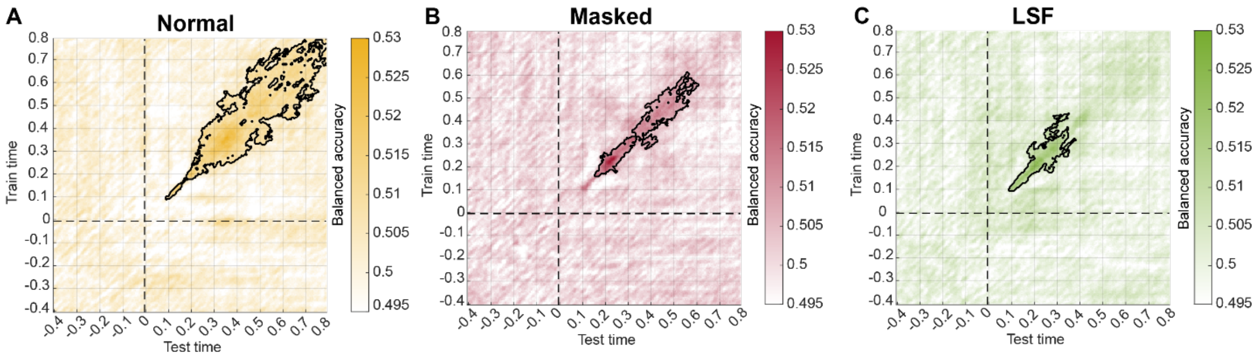


**Supplementary Figure 6: Temporal-generalization matrices for the superordinate category decoding analysis, corresponding to Fig. 3A-C (blue trace).** Matrices are shown for the (A) Normal, (B) Masked, and (C) LSF conditions. Colors indicate balanced accuracy, and the black outline indicates clusters that survived correction across training and testing time points ($p < 0.05$).

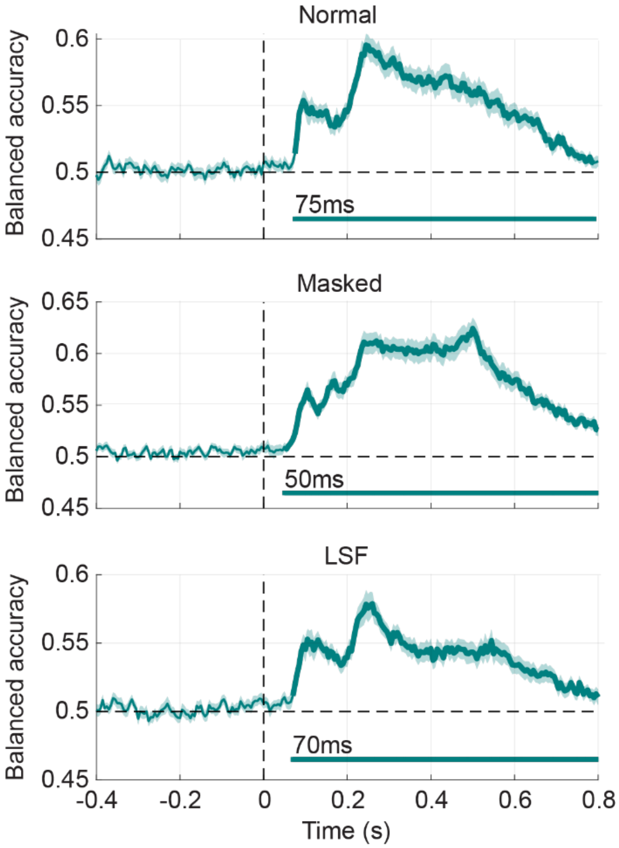


**Supplementary Figure 7: Decoding analysis results for position decoding in the Normal, Masked, and LSF conditions.** Shaded areas indicate s.e.m. across subjects (n = 30). Bolded traces and horizontal lines indicate significant timepoints ($p < 0.05$, cluster-based permutation test).

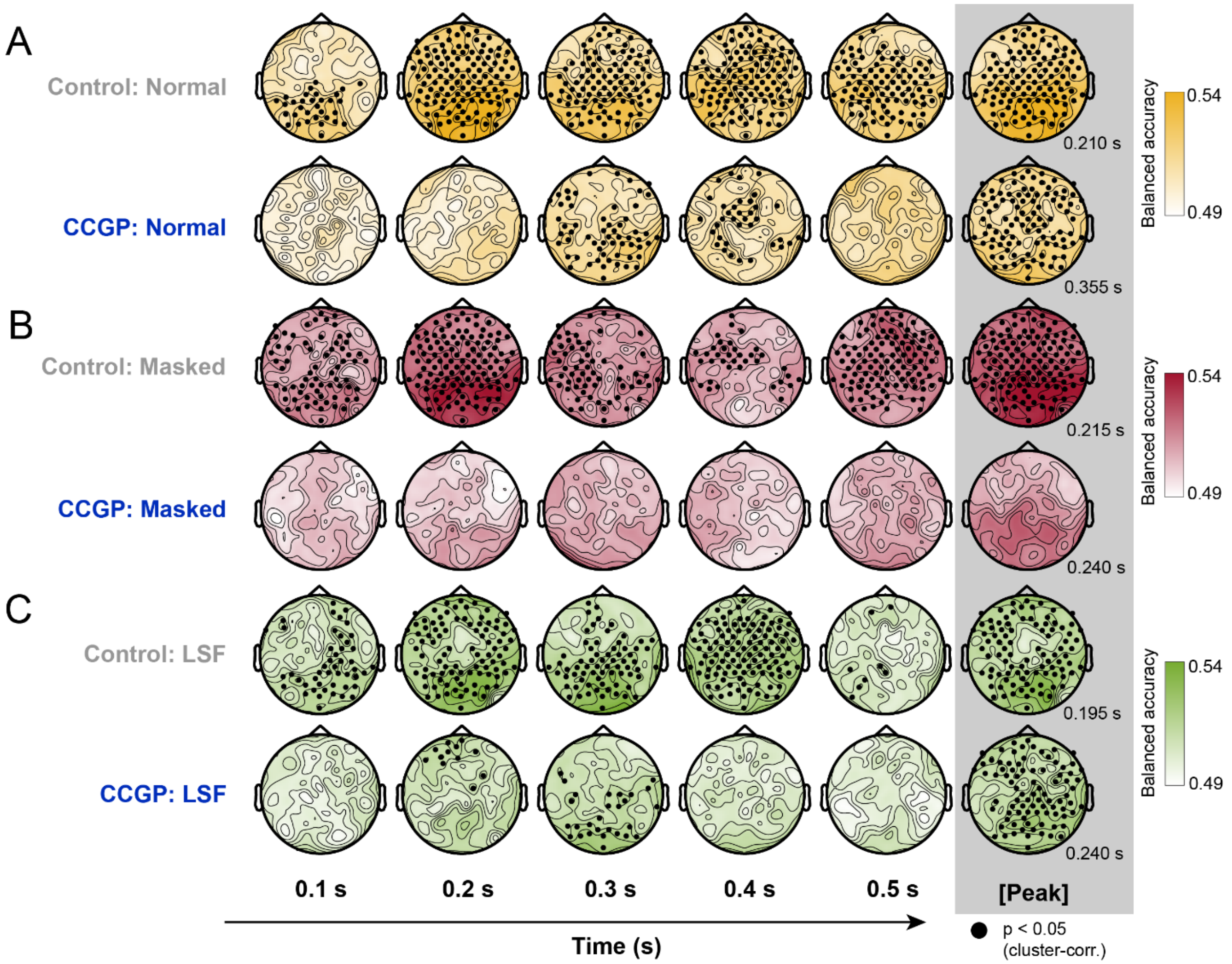


**Supplementary Figure 8: Searchlight decoding analysis results that complement the Cross-Category Generalization Performance (CCGP) and the corresponding control analysis in Fig. 4B-D.** See Figure 4A for the decoding analysis schematic. Top rows in each panel indicate the control decoding analysis, where classifiers were trained and tested within the same basic category pair, with a 90%-10% train-test split using a stratified cross-validation procedure. The bottom rows in each panel plot the CCGP analysis, in which classifiers are trained and tested on different basic category pairs (e.g., trained on dog vs. car and tested on cat vs. truck), yielding 4 decoding schemas that were averaged together. **(A-C)** Searchlight decoding analysis at the channel level from t = 0.1 s (100 ms after stimulus onset) to t = 0.5 s, and at peak decoding times (peaking decoding time indicated in the bottom-right corner) for **(A)** Normal, **(B)** Masked, and **(C)** LSF conditions. Black dots indicate significant channel clusters ($p < 0.05$, cluster-based permutation test across space and time).

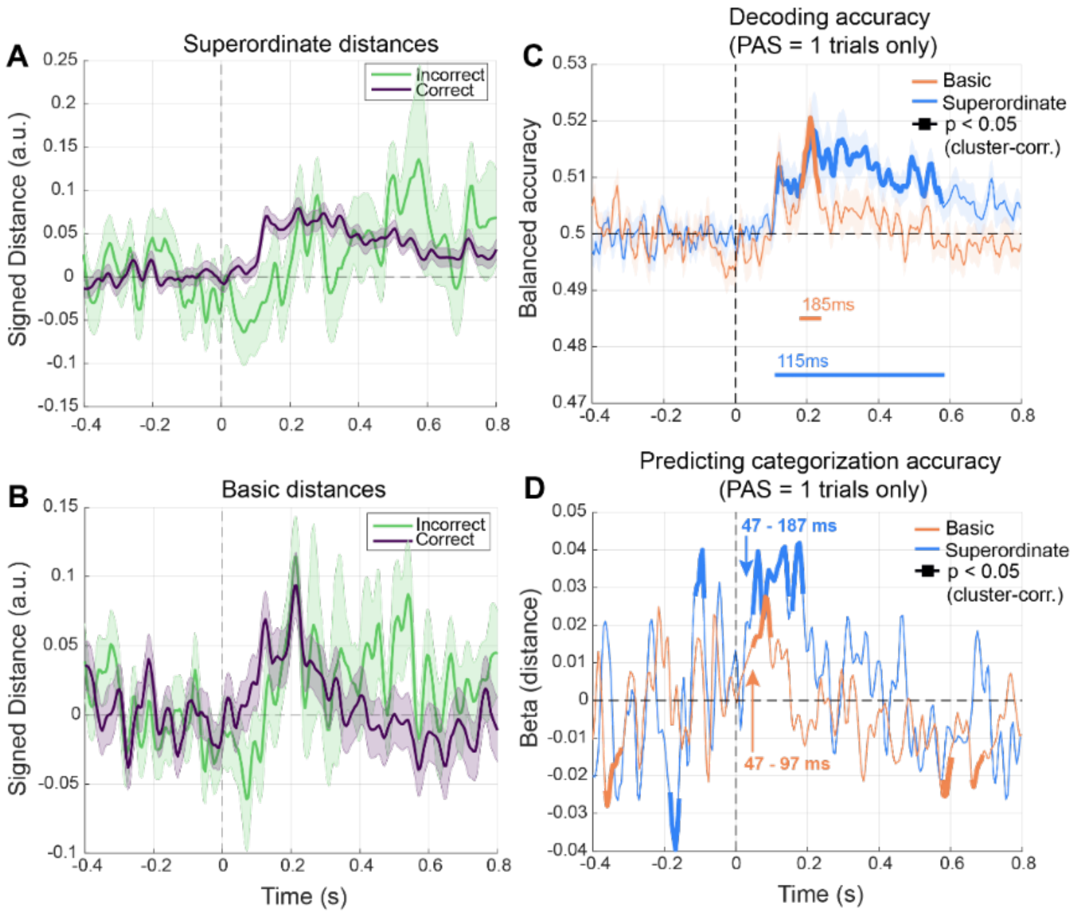


**Supplementary Fig 9: Distance-to-bound analysis testing the contribution of category representations to categorization behavior. (A)** Signed distance values for superordinate category representation, separated by correct or incorrect trials. Distance traces were smoothed using a Savitsky-Golay filter with a 150 ms window for visualization. **(B)** Same as (A), but for the basic-level category representation. **(C)** Decoding accuracy traces for the basic and superordinate level category decoding. Decoding analysis is similar to Fig. 3A-C but was pooled across the 3 conditions and restricting to trials with PAS rating reports of "1" (brief glimpse). In A-C, shaded area represents the s.e.m. across participants (n = 30). **(D)** $\beta_1$ values for the signed distance variable from logistic mixed-effects model, showing the relationship between trial-level signed distances and categorization performance ($logit(p_{i,k}) = \beta_1 N_{i,k,} + \sum_{c=2}^{3} \beta_{2,c} I(M_{i,k} = c) + b_{i)}$. See methods for details. In C-D, bolded traces indicate significant time points ($p < 0.05$; cluster-based permutation test). Numbers indicate the onset and offset times of the main significant clusters.
